# Plasma proteomic biomarkers identify non-responders and reveal biological insights about the tumor microenvironment in melanoma patients after PD1 blockade

**DOI:** 10.1101/2022.02.02.478819

**Authors:** Arnav Mehta, Marijana Rucevic, Emmett Sprecher, Milan Parikh, Jiajia Chen, Dennie T. Frederick, Elliot Woods, Lynn Bi, David Lieb, Lina Hultin-Rosenberg, Jamey Guess, Ryan J. Park, Alexis Schneider, William Michaud, Benchun Miao, Gyulnara Kasumova, Michelle S. Kim, Xue Bai, Russell W. Jenkins, Samuel J. Klempner, Anna L. K. Gonye, Keren Yizhak, Moshe Sade-Feldman, David Liu, Ryan J. Sullivan, Keith T. Flaherty, Nir Hacohen, Genevieve M. Boland

## Abstract

Most patients treated with immune checkpoint blockade (ICB) do not have durable treatment responses. Therefore, there is a critical need to identify early non-invasive biomarkers of response. We performed plasma proteomic analysis (>700 proteins) at three timepoints on 174 metastatic melanoma patients treated with ICB. We leverage independent training and testing cohorts to build a predictor of immunotherapy response that outperforms several tissue-based approaches. We found 217 differentially expressed proteins between ICB responders (R) and non-responders (NR), including a co-regulated module of proteins enriched in certain NR patients. By analyzing single-cell RNA-sequencing data of tumor biopsies from 32 patients, we dissected the relative contribution of cells in the tumor to proteins in circulation. The majority of proteins in the co-regulated NR module derived from tumor and myeloid cells. Amongst myeloid cells, we identified a subset of tumor-associated macrophages (TAMs) with a suppressive phenotype that expressed high levels of the co-regulated NR module, thus suggesting they are key drivers of non-response signatures. Together, our data demonstrates the utility of plasma proteomics in biomarker discovery and in understanding the biology of host response to tumors.

## Main

Immune checkpoint blockade (ICB) has revolutionized the treatment of many immunogenic tumors^1,2^ but a significant fraction of patients recur or do not respond to treatment, with median objective response rates (ORR) of 33-45% in metastatic melanoma patients on anti-PD1 (aPD1) monotherapy^1^. Early identification of non-responders is essential to mitigate unnecessary treatment toxicities and for alternative treatment approaches. However, predictive biomarkers for response are incompletely characterized and may be unique to patient subsets^3,4^. Several approaches have leveraged clinical variables and tissue biomarkers including tumor mutational burden (TMB), PD-L1 abundance, fraction of copy number alterations, HLA-I loss of heterozygosity (LOH), microsatellite status, interferon signatures, and immune composition from bulk RNA-sequencing^4–10^. While these have proven fruitful, the performance has limited clinical utility due to suboptimal predictive performance, the need for adequate tissue, and a failure to capture a global measure of host response.

Circulating biomarkers provide easy access for serial monitoring and can provide insight on the mechanisms of response to ICB^11–14^. In particular, peripheral blood analysis has revealed unique insight into the functional states and clonality of T cells after ICB^15–17^, but studies assessing systemic effects of ICB on host immune and tumor interactions are limited^18^. Importantly, changes in the plasma proteome in cancer patients treated with ICB are also poorly understood^19^. Two recent studies demonstrated a role of IL8 as a potential predictor of non-response to ICB^20,21^ and paired single-cell analysis from tumor and blood tissue suggests that myeloid cells may be a primary source for circulating IL8^21^. However, the use of plasma proteomics to reveal biological insight into immune responses has been limited by small sample sizes or targeted approaches focused on either individual or small subsets of cytokines^19,22^.

To identify circulating protein biomarkers of ICB responses, we used a proximity extension assay (PEA) to perform analysis of 707 proteins (**Table S1**) in 174 metastatic melanoma patients across three timepoints (baseline, 6 weeks on treatment, and 6 months on treatment) (**Fig. 1a, Extended Data Fig. 1**). ICB responders (R) were classified as those patients with disease control at 6 months including a complete response (CR), partial response (PR), or stable disease (SD); whereas ICB non-responders (NR) were classified as those who had progressive disease (PD) or requiring transition in therapy without radiographic response. A discovery cohort (cohort 1) consisted of 116 patients (66 R and 50 NR, clinical characteristics in **Table S2**) and an independent validation cohort (cohort 2) consisting of 56 patients (40 R and 14 NR). All exploratory analysis presented was performed using cohort 1. Cohort 2 was used strictly to validate findings.

**Figure 1.**
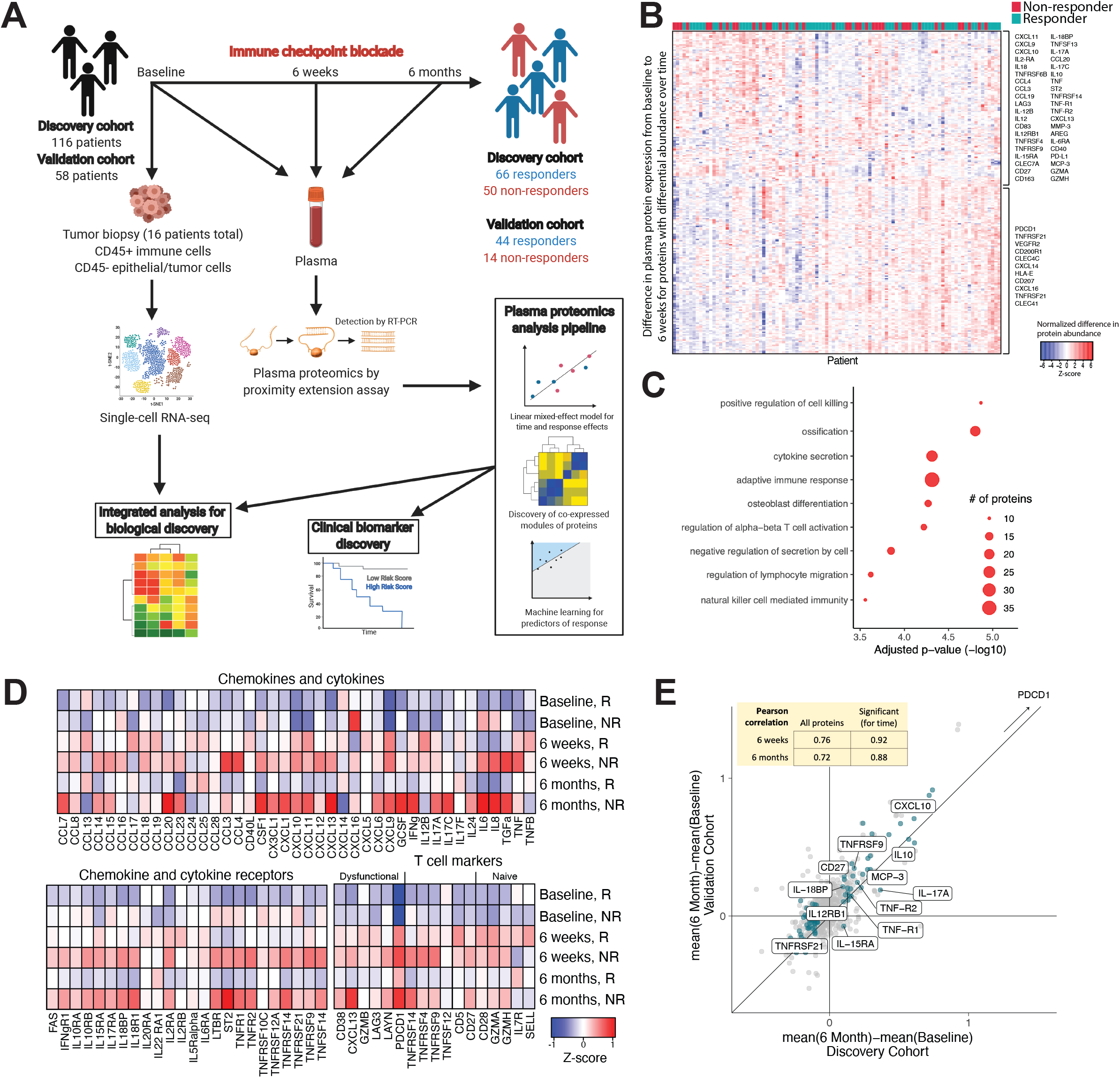
Melanoma patients treated with ICB have a unique treatment-associated pattern of inflammatory plasma proteins over time. (A) Schematic of our study design. (B) Heatmap showing difference in expression at 6-weeks post-ICB compared to baseline of differentially expressed plasma proteins over time (significant for time-effect in linear mixed-effect model). (C) Gene-ontology analysis for significant time-effect proteins. (D) Heatmaps demonstrating normalized mean expression of circulating cytokines, cytokine receptors and markers of T cell substates in plasma over time within treatment responder and non-responder subsets. (E) Scatter plot demonstrating correlation of plasma proteins in the discovery and validation cohorts. In green are proteins that were significant for time-effect in the discovery cohort.

Visualization of samples using principal-component analysis (PCA) showed that the samples did not clearly cluster by time point, tumor site, or BRAF status (**Extended Data Fig. 2**). To identify proteins significantly changed over time and between response groups, protein levels were fit to a linear mixed-effect model with main effects of time and of response, and an interaction effect of time and response (see **Methods**). In total, 247 proteins were identified as significant for time-effect (**Fig. 1b, Extended Data Fig. 3a and 4, and Table S3**) and an additional 162 for interaction effect (**Table S3**), including 98 overlapping proteins (**Fig. 2a**). When observing change from baseline to 6 weeks, proteins significant for time-effect separated into two distinct clusters by hierarchical clustering with generally opposing patterns of change (**Fig. 1b, Extended Data Fig. 4**). This was confirmed by co-abundance analysis of proteins across each timepoint. Time-effect proteins were enriched for those regulators of cytotoxicity, cytokine secretion, T cell and NK cell activation, and lymphocyte migration (**Fig. 1c**).

**Figure 2.**
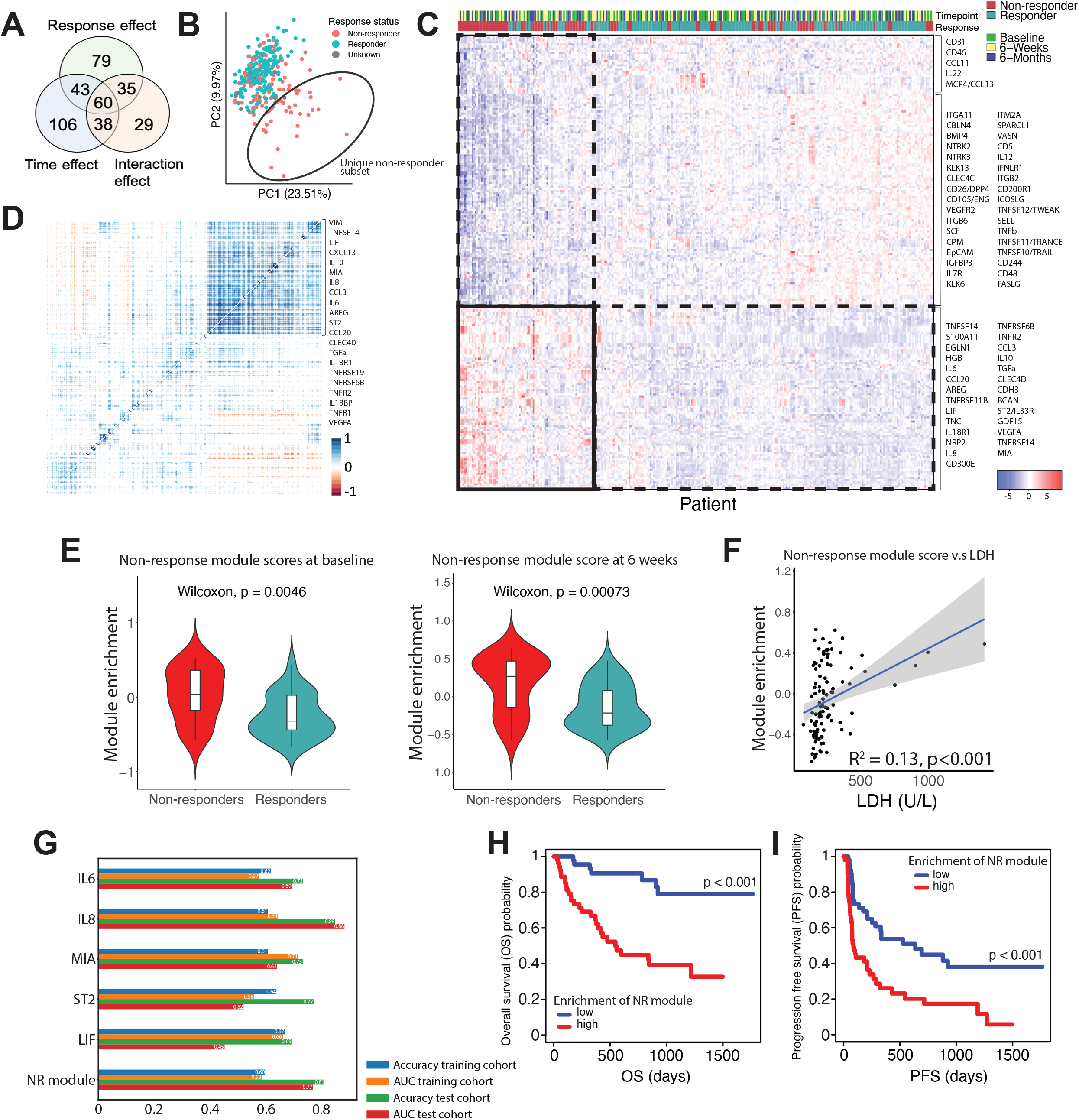
A co-varying module of plasma proteins is predictive of ICB non-response. (A) Venn diagram illustrating the total number of proteins and overlap between proteins significant for time, response and interaction effect in the linear mixed-effect model. (B) Principal component analysis of all patient samples color-coded by treatment response. The black oval represents a subset of non-responder samples that cluster separately from the rest of the samples. (C) Heatmap demonstrating normalized plasma expression for all proteins significant for response-effect in all patients and at all timepoints. (D) Heatmap demonstrating the correlation between plasma protein expression for all response-effect proteins. (E) Violin plots stratified by response status showing enrichment level of the non-response module set of proteins in the plasma of patients at baseline and at 6-weeks on treatment. (F) Scatter plot with a linear fit showing correlation of non-response module enrichment at baseline with LDH at baseline for each patient. (G) Accuracy in the discovery and validation cohorts, along with AUC in the validation cohort, of a logistic regression classifier for response status using each of the individual proteins shown and the non-response module enrichment score. (H)-(I) Kaplan-Meier survival curves for (H) overall survival and (I) progression free survival for patients stratified by expression of the non-response module enrichment score at baseline.

As previously demonstrated^19^, a dramatic increase in abundance of PD1 was observed following ICB treatment, as well as in several canonical proteins implicated in T cell recruitment and activity including CXCL9, CXCL10, CXCL11, GzmA, CD25 (IL2Ra) and 4-1BB (TNFRSF9) (**Fig. 1d,e and Extended Data Fig. 3a**). We detected several other cytokines, chemokines, and cytokine receptors (**Fig. 1d**). Whereas the fold-change on-treatment versus baseline for each protein varied, the vast majority of cytokines and cytokine receptors peaked at 6 weeks in Rs and at 6 months in NRs, with a notable normalization to baseline levels by 6 months in Rs (**FIg. 1d**). We detected several membrane-associated and intracellular protein markers of T cell substates, and found higher levels of markers of antigen-experienced and dysfunctional T cells (e.g. CD38, GzmB, LAG3 and LAYN) and those of naive T cells during treatment (e.g. CD28, IL7R and SELL (**Fig. 1d**)). Importantly, the time related changes correlated well with those seen in cohort 2 (pearson correlation of 0.92 at 6 weeks and 0.88 at 6 months) (**Fig. 1e**) confirming the reproducibility of these results.

To identify plasma proteins associated with immunotherapy response, we analyzed the 217 proteins significant for response-effect in our linear mixed-effect model (**Fig. 2a and Extended Data Fig. 3b**), which were validated as differential between R and NR in our validation cohort (**Extended Data Fig. 5;** R = 0.75 at 6 weeks and 0.77 at 6 months). PCA of all samples demonstrated that a subset of NR patients clustered separately from other patients and this was consistent at each individual time point (**Fig. 2b**). Hierarchical clustering of all samples by levels of response-effect proteins further highlighted the separation of a subset of NRs (**Fig. 2c**). Amongst the differentially abundant proteins were several cytokines and melanoma-associated proteins with reported associations to treatment non-response (**Fig. 2c,d and Extended Data Fig. 3b**), including IL6^23,24^, IL8^20,25,26^, MIA^26–28^, LIF^26,29^ and GDF-15^30^. These proteins were particularly abundant in the clustered subset of NRs (**Fig. 2b**) and were enriched for proteins regulating stress response, TNF signaling, leukocyte chemotaxis, and macrophage activation. Examination of cytokine and cytokine receptors suggested that the vast majority of inflammatory mediators peak at 6 weeks in R patients but continuously rise to 6 months in NR patients (**Fig. 1d and Extended Data Fig. 3b**). At baseline, ICB R patients notably had a trend towards higher levels of IFNg, CXCL11, and CCL13 (MCP4). CCL13 was the most significantly differentially expressed plasma protein with higher levels in Rs (**Fig. 1d and Extended Fig. 3b and 4b**).

Amongst plasma proteins higher in NR patients at baseline and 6 weeks were cytokines: IL6, IL8, TNFa, TGFa, CCL3 and CCL4 (**Fig. 1d, 2c and Extended Data Fig. 3b**). By 6 months these and several other cytokines were much higher expressed in NR patients, possibly reflecting a larger tumor burden at that time (**Fig. 1d**). The most significantly differential cytokine receptor was IL33R (ST2), more highly abundant in the plasma of NR (**Fig. 1d and Extended Data Fig. 3b**) and is highly expressed on tumor infiltrating regulatory T cells conferring poorer immunotherapy response^31,32^. To uncover which circulating plasma proteins may be co-regulated, we examined the covariance in levels of proteins significant for response effect in our linear mixed effect models (**Fig. 2d**). We uncovered a module of co-abundant proteins significantly more highly expressed in NRs (**Fig. 2d,e and Table S4**; herein referred to as the “non-response module”) at baseline and 6 weeks (**Fig. 2e**). We found that LDH (proxy for tumor burden), though significantly correlated, accounts for less than 13% of the variance in non-response module enrichment scores (**Fig. 2f**), implicating that any heterogeneity in this module can not be explained by tumor bulk alone. While most proteins were not correlated with tumor bulk, a select few (e.g. IL8) appeared to correlate (**Extended Data Fig. 6**).

To investigate if individual proteins in the non-response module may be predictive of immunotherapy response, we used logistic regression classifiers for each protein to predict response status in our discovery cohort (training cohort), and tested the accuracy of these classifiers in our discovery cohort (test cohort) (**Fig. 2g**). We found overall that classifiers trained on the difference in protein levels from baseline to 6-weeks performed better than levels at either baseline or 6-weeks alone. IL8 had the best predictive performance overall (AUC 0.88 in validation cohort), however, other proteins previously implicated in melanoma immunotherapy resistance performed poorly (IL6: AUC 0.69; MIA: AUC 0.64; ST2: AUC 0.52; LIF: AUC 0.45) compared to enrichment of the non-response module (AUC 0.77) (**Fig.2g**). Other machine learning methods using the entire protein dataset demonstrated variable performance with an AdaBoost classifier performing best overall (AUC 0.80), compared to a random forest classifier (AUC 0.70), parameter optimized k-nearest neighbor classifier (AUC 0.71), neural network (multi-layer perceptron; AUC 0.68), naive Bayes classifier (AUC 0.54) and logistic regression classifier (AUC 0.75) (**Extended Data Fig. 7**). Of note, almost all of these classification approaches performed less well to logistic regression on our non-response module score (**Fig. 2g**).

To assess the impact of individual protein levels versus the non-response module on survival, we stratified patients as high or low expressors by median levels at baseline and performed survival analysis using Kaplan-Meier curves with overall survival (OS) and progression free survival (PFS) (**Fig. 2h,i and Extended Data Fig. 8**). We found that enrichment of our non-response module strongly stratified patients with poor OS (**Fig. 2h**; p<0.001) and PFS (**Fig. 2i**; p<0.001). Several individual proteins were also prognostic of poor OS, including IL8, ST2, MIA and LIF (**Extended Data Fig. 8**). Of note, CCL13 (MCP4) was the only protein at baseline that conferred a better OS, but did not significantly predict investigator-assessed PFS.

We next leveraged scRNAseq data from tumor immune cells of patients in this cohort^33^ (n=16) and from malignant cells of patients in an independent cohort^34^ (n=14) to identify cell subsets and substates that express mRNA for proteins that are differential in the plasma of ICB R and NR patients (**Fig. 3 and Extended Data Fig. 9-16**). In parallel analysis of tumors and plasma from our patients, we found that a subset of genes had concordant changes in expression between R and NR patients, and correlated between plasma protein abundance and tumor gene expression (**Extended Data Fig. 12**; overall R=0.36 for all response effect proteins/genes). This suggested that a subset of response effect proteins in plasma may reflect changes within the TME during immunotherapy.

**Figure 3.**
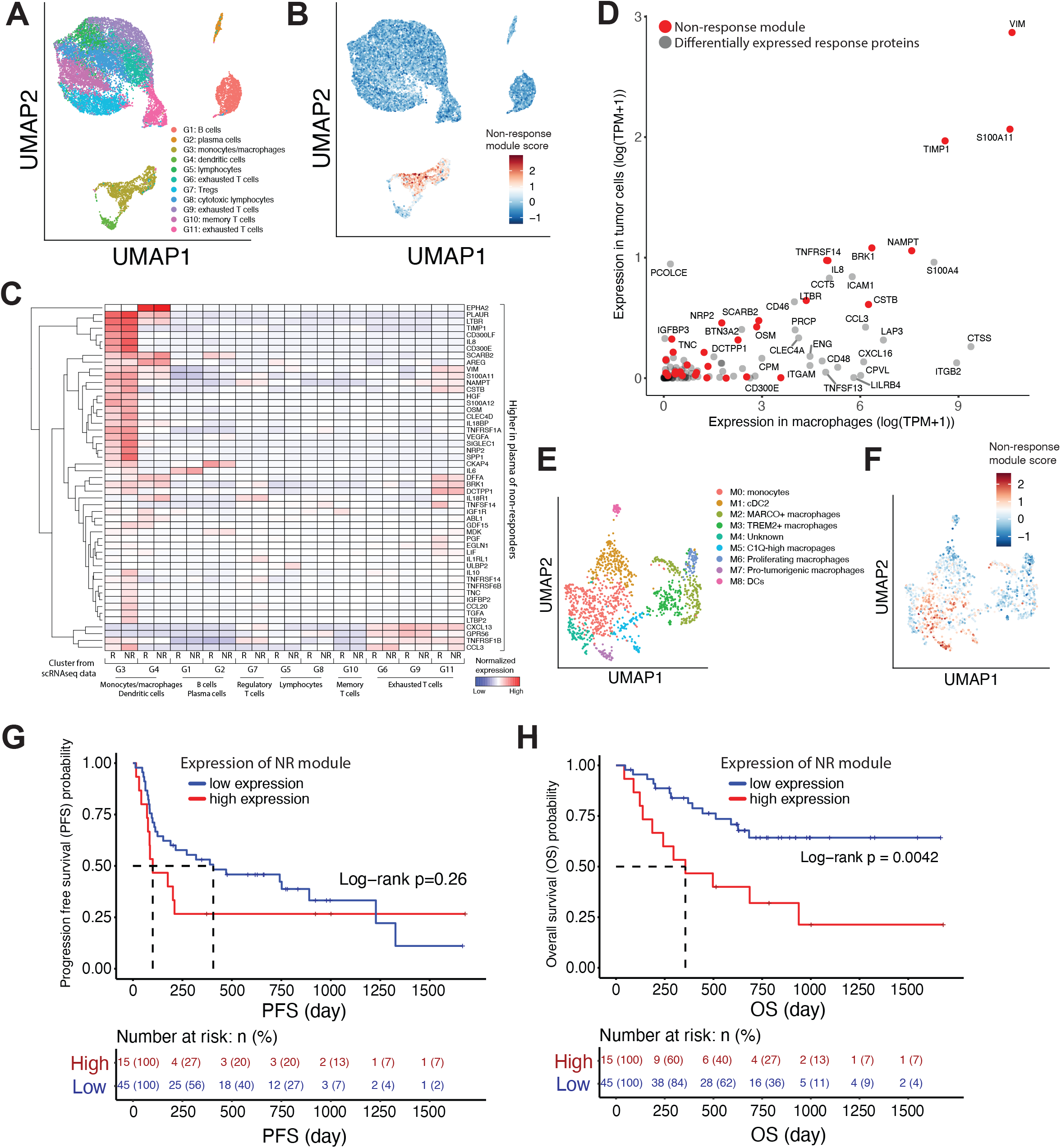
A suppressive tumor macrophage subset and tumor epithelial cells have enriched gene expression for the plasma-derived non-response module of proteins. (A) UMAP embedding of immune cell scRNAseq data from Sade-Feldman et al.^33^ with annotation of major cell types and subsets. (B) Non-response module enrichment in individual immune cells from (A) visualized on the UMAP embedding. (C) Heatmap showing pseudobulked expression of all plasma proteins elevated in the plasma of NRs within immune cell subsets in (A) separated by cells found in immunotherapy Rs versus NRs as previously annotated^33^. (D) Scatter plot showing gene expression counts as log(TPM+1), for all response-effect proteins discovered in plasma, in tumor epithelial cells versus tumor macrophages within the TME. Data represented is obtained by scRNAseq analysis of biopsies obtained from patients in this plasma proteomic cohort either from previously generated immune cell subsets^33^ or newly generated tumor epithelial cells. (E)-(F) Subclustering analysis of macrophages in (B) visualized in a UMAP embedding and showing (E) annotated macrophage subsets and (F) non-response module enrichment score within these cells. (G) Progression free survival (PFS) and (H) overall survival stratified by high versus low non-response module score (split by quartile; n high = 15, n low = 45) in a subset of patients from Schadendorf cohort^8^ with relatively high myeloid proportions (n= 60). Tumors with high non-response module had worse OS (two-sided KM log-rank test, P = 0.0042), with consistent trend in PFS (two-sided KM log-rank test, n.s.).

We next evaluated the enrichment of our non-response module with individual subsets of immune cells in the TME (**Fig. 3a,b and Extended Data Fig. 9a**). Unsupervised clustering of all cells revealed eleven distinct clusters of immune cells as previously described (**Fig. 3a**). We found that the non-response module was most enriched in monocytes and macrophages (**Fig. 3b and Extended Data Fig. 9a, 13**; cluster G3), and we verified these findings in an independent scRNAseq melanoma dataset^34^ (**Extended Data Fig. 16**). To identify specific proteins contributing to the expression of the non-response module in tumor myeloid cells, we examined pseudobulked gene expression of all response effect proteins in each subset of immune cells separated by response status (**Fig. 3c and Extended Data Fig. 10**). Transcripts corresponding to proteins associated with non-response were more highly expressed in tumor-infiltrating immune cells compared to response proteins (**Fig. 3c and Extended Data Fig. 10**). To determine if circulating myeloid cells contribute to plasma proteins in our non-response module, we assessed the correlation between plasma non-response module score or individual plasma protein abundance with absolute monocyte count (AMC) or absolute neutrophil count (ANC). We found overall that myeloid cell counts did not account for the variance in plasma levels of the non-response module and individual plasma proteins in this module (**Extended Data Fig. 9b, 11**), thus suggesting that abundance of circulating myeloid cells alone does not account for enrichment of the non-response module.

To investigate the contribution of melanoma tumor cells to circulating non-response proteins, we analyzed scRNAseq data from tumor epithelial cells from 21 patients, including 7 patients in this cohort^34,35^ (**Extended Data Fig. 14, 16**). We found that the enrichment of the non-response module was heterogeneous within tumor cells (**Extended Data Fig. 14c, 16e-g**), with some enrichment of module expression in particular epithelial cell subsets. Genes expressed uniquely in tumors were canonical melanoma-associated genes such as MIA (melanoma inhibitory antigen), known pro-tumorigenic proteins such as TNFRSF19, and putative novel regulators of resistance with strong differential signatures including BCAN, CDH3, MFAP5, and SERPINA5 (**Extended Data Fig. 14c, 16e-g**).

Given the contribution of both tumor myeloid and epithelial cells to levels of the plasma non-response module, we dissected the relative contribution of each cell population to plasma proteins. Analysis of the pseudobulked expression of all cells within each subset in both scRNAseq datasets revealed that a large subset of proteins in the non-response module had high expression in both tumor cells and macrophages (**Fig. 3d and Extended Data Fig. 17,18**), including VIM, S100A11, TIMP1, BRK1, NAMPT and IL8 (**Fig. 3d**). Importantly, we found several proteins in the non-response module (CD300E) and otherwise associated with non-response (CTSS, ITGB2, TNFSF13, CD48, CXCL16, among others) that were uniquely expressed in macrophages, and several known and putative novel melanoma resistance proteins in the non-response module (BCAN, MIA, CDH3, NRP2, LTBR, among others) uniquely expressed within tumor epithelial cells (**Fig. 3d and Extended Data Fig. 17,18**).

To identify macrophage subsets driving expression of the non-response module, we performed a subclustering analysis of monocytes and macrophages from the scRNAseq data of patients in our cohort (**Fig. 3e,f and Extended Data Fig. 15**) and determined marker genes for each cluster (**Table S5**). We found one particular macrophage subset (cluster M7 in **Fig. 3e**) that had the highest expression of the non-response module (**Fig. 3f and Extended Data Fig. 15c,d**). This subset, which we term pro-tumorigenic macrophages and expresses PCOCE2 and SIGLEC1 among top marker genes, closely resembled tissue resident tumor-associated macrophage subsets recently described^36,37^ that establish a niche favorable for tumor growth. Three other subsets (clusters M0, M4 and M5) demonsted heterogeneous expression (**Extended Data Fig. 15c,d**) of the non-response module and were composed of monocytes and C1Q^high^ macrophages as previously described^38^.

This suggests that the M7 macrophage subset, in addition to it’s reported pro-tumorigenic properties, may also be a poor prognostic sign for immunotherapy response, and may contribute to the abundance of circulating non-response associated proteins in some patients. To validate the prognostic significance of the non-response module in macrophage populations, we analyzed bulk RNA-seq data from an independent cohort of melanoma patients^8^. We focused on tumors with a high myeloid cell fraction to perform a survival analysis, and found that tumors that scored highly for the non-response module signature had overall poorer prognosis, with lower OS (**Fig. 3g,h and Extended Data Fig. 19**).

In summary, we leverage an integrated analysis of plasma proteomics across serial timepoints and tissue scRNAseq in melanoma patients treated with immune checkpoint blockade to shed insight into the biology of immunotherapy non-response. While previous studies focused on individual protein biomarkers (e.g. IL8) or small cohorts without paired timepoints, our high-plex proteomic analysis in 174 patients with serial monitoring enabled the characterization of two distinct subset of immunotherapy non-responsive patients and a co-varying group of plasma proteins (referred to as the non-response module) that is enriched in NRs. We find this non-response module is strongly predictive of which patients will respond to immunotherapy, and outperforms several machine learning approaches trained on the entire protein dataset and other published prediction approaches using only tissue-based features^8,9,39,40^. A shortcoming of our predictive approach is that we can reliably predict responders, and hence have a high positive predictive value, but are able to less effectively predict NRs due to the heterogeneity in plasma profiles of non-responder patients, and hence have a low negative predictive value. Clinically, this is favorable as we aim to ensure no patient that might benefit goes untreated. However, this highlights a key need to better endotype non-responder patients and the importance of future work leveraging both plasma and tumor features for maximum clinical impact.

We noted the presence of myeloid associated proteins IL8^20,21^, VEGFA^41^ and CLEC4D^42^, the Treg proteins TNFR2^43^ and AREG^44^, the IL-18 sink IL-18 binding protein (IL-18BP)^42,45^ and IL-10^46^, which has pleiotropic effects on immune cells in the non-response module. Our findings are largely consistent with previous work describing IL8 as a potential negative prognostic biomarker of immunotherapy response^20,21^. Whereas these studies correlated abundance of circulating IL8 with peripheral neutrophil/monocyte counts and tumor CXCL8 expression, our unbiased approach enabled a deeper understanding of co-varying sets of proteins with elevated levels with IL8 in circulation. This enabled a more refined understanding of tumor and immune cell subsets in the TME that may confer poor overall outcome.

While interpretation of the source of circulating plasma proteins may be confounded, we identify tumor cells and a subset of pro-tumorigenic macrophages as likely primary contributors to enrichment of the non-response module in plasma. Our findings are consistent with recent literature suggestive of tissue resident tumor-associated macrophages that interact with tumor epithelial cells and promote tumor growth. In particular, we note a strong correlation in gene expression signatures between our identified M7 populations, which have the strongest enrichment of the non-response module, and the IL4l1+ pro-tumorigenic macrophage subset (cluster 6) in Mulder et al^38^. Consistently, we find our cluster M7 has strong overlapping marker features with the Group I macrophages as described in Casanova-Acebes et al.^36^ Cluster M0, which has weaker enrichment of the NR module, likely represents the progenitor ISG high monocyte population of IL4l1+ macrophages as identified in cluster 4 of Mulder et al^38^. Finally we note strongly correlated expression signatures between our cluster M5, which also showed some enrichment of the NR module, and the HES1, C1Q-high and TREM2 macrophage populations identified in Mulder et al^38^. Whether these macrophage subsets are implicated in establishing an immunotherapy-resistant niche or have specific immunosuppressive function after treatment remains to be explored and will need to be evaluated in larger cohorts with serial tissue sampling. Importantly, the contribution of peripheral and other tissue myeloid cells towards plasma enrichment of the non-response module needs to be further elucidated. Importantly, it remains unclear how enrichment of the non-response module may relate to T cell phenotypes and clonality^15,40,47^ in the setting of ICB, which will also need to be explored in larger paired plasma and tissue cohorts.

## Acknowledgements

We would like to thank all the patients and the families that participated in our biobanking effort and enabled this study using plasma samples. We thank Olink Proteomics for their in-kind contribution to process all plasma samples for proteomics in this study. We also thank the MGH Cancer Center, Department of Medicine and Department of Surgical Oncology for enabling the proposed studies. This work was supported by the Massachusetts General Hospital Jackson Society White Coat Grant (AM), the NIH 5T32CA071345-23 (AM), a BMS Merit Award (XB), NIH K08CA226391 (RWJ), the Melanoma Research Alliance Young Investigator Award (RWJ, DL), Doris Duke Charitable Foundation Clinical Scientist Development Award (DL), NIH K08CA234458 (DL), the Termeer Early Career Fellowship in Systems Pharmacology (RWJ), NIH R01 CA229851 (RJS), Adelson Medical Research Foundation (KTF, NH), NIH/NCI U54 CA224068 (NH), NIH/NCI R01CA208756 (NH) and a chair and gift from Sandra, Sarah and Arthur Irving (NH).

## Author contributions

Conceptualization: AM, MR, RJS, KTF, NH, GMB. Resources: AM, MR, NH, GMB. Methodology: AM, MR, ES, RWJ, SK, DL, RJS, KTF, NH, GMB. Data generation and clinical annotation: AM, MR, DTF, EW, WM, BM, GK, MSK, XB, ALKG, KY, MSF. Formal analysis: AM, MR, ES, MP, JC, DTF, EW, LB, DL, LHR, JG, RJP, AS. Writing – Original Draft: AM, NH, GMB. Writing – Review and Editing: AM, MR, RWJ, SJK, DL, RJS, KTF, NH, GMB with contributions from all authors.

## Methods

### Collection of patient plasma samples

Metastatic melanoma patients at MGH provided written informed consent for the collection of blood samples (DF/HCC IRB approved Protocol 11-181). Whole blood was collected in BD Vacutainer CPT tubes (BD362753) prior to treatment and 6 weeks on treatment with immune checkpoint blockade, and 3mL of plasma was isolated after centrifuging tubes for 30 minutes at room temperature. Plasma was stored at -80 ºC for further use and thawed at the time of proteomic analysis.

### Plasma proteomic immunoprofiling

A highly multiplexed Proximity Extension Assay (PEA) was applied to measure protein abundance directly in the plasma obtained from metastatic melanoma patients. PEA is an affinity-based assay developed by Olink Proteomics as previously described^48^. In brief, for each protein, a pair of oligonucleotide-labeled antibody probes bind to the targeted protein and if the two probes are in close proximity, a PCR target sequence is formed by a proximity-dependent DNA polymerization event and the resulting sequence is subsequently detected and quantified by real-time PCR using the Fluidigm BioMark™ HD real-time PCR platform. Here, the abundance levels of 736 plasma proteins were analyzed in 347 metastatic melanoma specimens collected from 116 patients at the baseline, 6-weeks and 6-months using 8 pre-defined PEA-panels (the Olink Multiplex Inflammation, Immune Response, Immuno-Oncology, Oncology II, Oncology III, Cardiometabolic, Cardiology III and Neurology) and quantified by real-time PCR. A complete list of the 736 assays corresponding to 630 unique proteins are listed in **Table S1**. The patients’ plasma samples were randomized across four 96-well plates such that each sample was assayed once and normalized for any plate effects using the built-in inter-plate controls according to Olink’s recommendations. The resulting abundance levels are given in NPX (Normalized Protein eXpression) that is on log2-scale. Each PEA has an experimentally determined lower limit of detection (LOD) calculated based on negative controls that are included in each run and measurements below this limit were removed from further analysis. Assay characteristics including detection limits, assay performance and validations are transparently available at www.olink.com. Subsequently, in the same manner we have analyzed the levels of 978 plasma proteins in melanoma specimens obtained from validation cohort (n=55 patients) at the baseline, 6-weeks and 6-months using 11 pre-defined PEA-panels (the Olink Multiplex Inflammation, Immune Response, Immuno-Oncology, Cell Regulation, Oncology II, Metabolic, Cardiometabolic, Cardiology III, Cardiology II, Neurology, Organ Damage).

### Pre-processing of plasma proteomic data

All analysis of plasma proteomic data was performed in the R (version 4.0.3) programming environment unless otherwise specified. A data frame of plasma proteomic data in the form of NPX values was loaded into R using the OlinkAnalyze package (https://github.com/Olink-Proteomics/OlinkRPackage). Of the 736 assays in total, 29 assays were found to have greater than 75% of samples below the LOD, and were excluded from further analysis. Of the 347 metastatic melanoma specimens in the discovery cohort, 18 samples failed quality control in at least one Olink panel and were subsequently excluded from further analysis. In addition, 9 patients were additionally excluded because they had received confounding therapies, thus leaving a total of 321 samples in the discovery cohort, including 108 baseline samples, 113 6-week samples and 100 6-month samples. The final validation cohort consisted of 119 samples, including 46 baseline samples and 26 6-week samples, and 1043 assays after similar filtering steps.

### Dimensionality reduction and visualization of proteomic data

Principal components analysis was performed on the plasma proteomic data for dimensionality reduction using the prcomp function in the prcomp package in R. Principal components were visualized using the autoplot function in the prcomp package in R. Non-linear embeddings were performed for visualization using Uniform Manifold Approximation and Projection (UMAP) using the umap function in the umap package in R. UMAP embeddings were visualized using a custom built plotting function that is available in the Github repository for this paper as described below. All heatmaps in the paper were generated using the heatmap.2 or heatmap3 function in R.

### Linear mixed effect models and mean different estimates

Linear mixed effects models were fit independently to each assay using the lme4 package in R. The models included a main effect of time, a main effect of responder status, the interaction between time and responder status and a random effect of patient to account for the correlation between measurements coming from the same individual. Significance of the three model terms was determined with an F-test using Satterthwaite degrees of freedom and implemented with the lmerTest package in R. All p-values were adjusted to control the false discovery rate (FDR) at 5% using the Benjamini-Hochberg method. This approach was applied separately to the discovery cohort and the validation cohort.

Estimated differences in NPX means between responders and non-responders were calculated from the linear mixed effects models for each assay using the emmeans package in R. P-values for estimates from each assay were adjusted using the Tukey method. The same approach was used to estimate differences between mean NPX at week 6 compared to baseline and month 6 compared to baseline. All code and custom functions used in this analysis are available in the paper’s Github repository as described below. Results from this analysis are in **Table S3**.

### Correlation scatterplots

Pearson correlations of estimated mean differences (1) between responders and non-responders at each timepoint and (2) from baseline to either 6-weeks or 6-months were calculated between the discovery and validation cohorts using the cor function in R. The estimates for each cohort were also plotted in a scatterplot using the ggplot package in R. Assays were colored as statistically significant if the main effect of response or the interaction effect for response and time passed the FDR threshold for the F-test and the Tukey adjusted p-value for the estimated mean difference was less than 0.05.

### Module score for each plasma proteomic sample

Samples were scored for sets of genes, including the non-response module, using single sample gene-set variance analysis implemented in the GSVA package in R. Non-response module enrichment was subsequently plotted using the ggplot package in R.

### Survival analysis

Survival analysis was performed using the survfit function in R for either overall survival or progression-free survival after subsetting patients into high expressors and low expressors by median expression of each protein of interest. A log-rank test statistical test was performed using the survdiff function in R to calculate p-values for survival curves with the null hypothesis that there was no difference in survival between the high and low expressing patient subsets for each protein.

### Prediction of immunotherapy response

All analysis for predictions was done in Python using NumPy, SciPy, Matplotlib, and scikit-learn. Machine learning methods were trained on normalized protein expression (NPX) matrices from a training cohort of 116 patients and tested on an independent validation cohort of 58 patients. NPX matrices were filtered to include only patients with a response classification (responder, R, and non-responder, NR; 4 patients had unknown response), and patients with both baseline and 6 week samples (107 patients in the training cohort and 26 patients in the validation cohort). A larger panel of 1040 assays was used in the validation cohort, so the count matrix was filtered to only those assays overlapping the training cohort (598 assays total). Separate count matrices were then created with NPX values at baseline, 6-weeks, or the difference from baseline to 6-week (delta NPX) for each cohort.

A selection of classification models was chosen from scikit-learn including: Random Forest, K Nearest Neighbors (KNN), Multi-layer Perceptron Classifier (MLP), Gaussian Naïve Bayes, AdaBoostClassifier (based on the Decision Tree Classifier), and Logistic Regression (LogReg). The best hyperparameters for each model were found using grid searches with 4-fold cross-validation on the training dataset. Each model was then trained on the NPX values with these hyperparameters and tested for accuracy in predicting patient response against both the training and validation datasets. ROC curves and AUC were calculated for each (sklearn.metrics.roc_curve, sklearn.metrics.roc_auc_score) against the validation dataset. This process was repeated for the baseline, 6-week, and delta NPX matrices, and it was found that the delta matrix was most predictive of response across all models. We found that the KNN, LogReg, and MLP classifiers performed best on the validation cohort. Several individual proteins that were most highly differentially expressed between responders and non-responders (e.g. IL6, IL8, MIA, ST2, LIF) were tested using the Logistic Regression model with the baseline, 6-week, and delta matrices. Accuracy against the training and validation datasets, ROC curves, and AUC were calculated for each as above.

### Generation of plate-based single-cell RNA-sequencing data from tumor samples

For the generation of scRNAseq data from tumor epithelial cells for patients in this cohort, we closely followed our protocol as previously described^33^. Briefly, freshly isolated tumor samples were dissociated using the human tumor dissociation kit (Miltenyi Biotec; 130-095-929): surgical resection tissue was first minced into small pieces using a scalpel, and placed in a 1.5mL eppendorf tube containing enzyme H (100uL), enzyme R (50uL), enzyme A (12.5uL) and RPMI (837.5uL). The sample was incubated for 20 minutes in a thermomixer at 37degC at 600rpm. The sample was subsequently strained with a 70uM cell strainer and washed with cold PBS containing 1.5% FBS. Samples were subsequently spun down at 1300rpm in 4degC for 5 minutes, and resuspended in PBS for cell counts and to assess viability with trypan blue using a Countess automated cell counter (Invitrogen). To sort CD45+ cells, samples were stained with Human TrueStain FcX (Biolegend; 422303) and Zombie Dye Violet Blue (Biolgend; 423114) for 15 minutes at 4degC followed by surface labeling with PE anti-human CD45 (Biolegend; 304008) for 30 minutes at 4degC. CD45-cells were then sorted into 96-well plates (Eppendorf; 951020401) containing 10uL of lysis buffer (TCL buffer, Qiagen 1031576 supplemented with 1% beta-mercaptoethanol), sealed, vortexed, spun down at 3500rpm for 30 seconds, and immediately placed on dry ice and stored at -80degC. At the time of processing, samples were thawed on ice and processed using the previously described Smart-Seq2 protocol^33^ for single cell libraries. Combined libraries for 384 cells were sequenced on an Illumina NextSeq 500 sequencer using paired end 38-base pair reads.

### Analysis of SmartSeq2 single-cell RNA-sequencing data

Single-cell RNA-sequencing data was either generated as described above or obtained as described from previous studies^33–35^. Demultiplexed FASTQ files obtained from sequencing were aligned using STAR^49^ to NCBI Human Reference Genome GRCh37 (hg19). Per-cell gene expression levels were quantified using transcripts per million (TPM) using RSEM^50^. All downstream single-cell analysis was performed in the Seurat (version 4.0.4) package in R. Briefly, raw transcripts per million (TPM) count matrices were log1p transformed as log(TPM+1). Highly variable genes were selected using the Seurat FindVariableFeatures() with default parameters. Dimensionality reduction was subsequently performed either for the entire dataset or for particular subsets of cells using the Seurat RunPCA() function with default parameters. The top 40 PCs were used to generate a k-nearest neighbor graph using the Seurat FindNeighbors() function and clusters were identified using shared nearest neighbor (SNN) modularity optimization with the Seurat FindClusters() function. UMAP embeddings were calculated using the Seurat RunUMAP() function and visualized using the Seurat DimPlot() or FeaturePlot() functions. For cluster marker genes, the Seurat FindAllMarkers() function was used. Cells were scored for expression of the non-response module using the Seurat AddModuleScore() function, which finds the average expression level for non-response module genes and subtracts from it the expression of a control set of genes that have the same binned mean expression levels across the entire dataset. Cells were further scored for cluster signatures from Mulder et al^38^ and Casanova-Acebes et al.^36^ using the AddModuleScore() function using the 100 most significant differential marker genes represented in our dataset for each cluster. Cluster scores were correlated using the corr() function in R, and plotted using the corrplot() function. Log normalized count matrices were used to generate scatter plots of pseudobulked expression of different immune subsets using the ggplot function in R.

### Survival analysis using bulk RNA-sequencing data

Patients in Schadendorf cohort had advanced melanoma and had received PD-1 immune checkpoint blockade (ICB)^8^. 121 out of 144 patients had bulk RNA-seq data available, among which 56 patients with progressed disease (PD) as best response to anti-PD-1 ICB were identified as progressors, 47 patients with complete response (CR) or partial response (PR) were identified as responders. Remaining 2 patients with mixed response (MR) or 16 stable disease (SD) were included for deconvolution analysis and assessment of survival outcome but were excluded from correlational analysis.

Deconvolution was performed with a gene signature of myeloid population derived from scRNA-seq. The myeloid scores were calculated using gene set variation analysis in the R package GSVA. The NR module score for each sample was calculated using the mean of z scores of 68 genes from non-response module. Correlation between myeloid scores and non-response module scores was computed with nonparametric Spearman’s rank correlation coefficient.

A subset of patients (n=60) with high myeloid proportion (split by median of myeloid scores) were included in survival outcome evaluation. In Kaplan-Meier survival analysis, the significance of the difference in survival outcome between the patients with high non-response module score (top quartile) and low non-response module score (bottom quartile) was assessed using a two-sided log-rank test.

### Data Availability and Code

All plasma proteomic data will be made publicly available at the time of publication. All code and Jupyter notebooks associated with the analysis of the plasma proteomic and single-cell RNA-seq data will be made available here: https://github.com/arnav-mehta/melanoma_proteomics. The authors will provide any further detail as necessary.

## Extended Data Figure legends

**Extended Data Figure 1**. Schematic of the distribution of timepoints for each patient sample. Labeled on the y-axis is patient ID, and the line plots indicate the timepoint for collection for baseline, 6-week and 6-month samples for all samples in the discovery cohort.

**Extended Data Figure 2**. (A)-(B) Principal component analysis of all patient samples color-coded by (A) treatment timepoint and (B) site of primary lesion; shown is a plot of PC1 v.s. PC2.

**Extended Data Figure 3**. (A)-(B) Heatmaps of mean plasma protein expression for proteins significantly differentially expressed over (A) time and (B) response status shown stratified by timepoint and response status.

**Extended Data Figure 4**. (A) Heatmap demonstrating normalized plasma expression for all proteins significantly differentially expressed over time in all patients and at all timepoints. (B) Point-range plots split between responders and non-responders of CCL13 (MCP-4), IFN-gamma and CXCL11.

**Extended Data Figure 5**. Scatter plot demonstrating correlation of mean difference in plasma proteins between treatment Rs and NRs in the discovery and validation cohorts. In green are proteins that are differentially expressed by response status.

**Extended Data Figure 6**. Scatter plot and linear fit showing normalized protein expression for IL6, IL8, LIF or MIA versus LDH for each sample.

**Extended Data Figure 7**. Accuracy in the discovery and validation cohorts, along with AUC in the validation cohort, of several machine learning models trained on the entire protein set as features.

**Extended Data Figure 8**. Kaplan-Meier survival curves for overall survival for patients stratified by median baseline expression of IL8, ST2, MIA or LIF.

**Extended Data Figure 9**. (A) Histograms of non-response module enrichment scores in cells within each of the immune cell clusters described in Fig. 3b. (B) Correlation of non-response module enrichment score and absolute monocyte count (AMC) or absolute neutrophil count (ANC) for each sample.

**Extended Data Figure 10**. Heatmap showing pseudobulked expression of all plasma proteins elevated in the plasma of Rs within immune cell subsets in Fig. 3b separated by cells found in immunotherapy Rs versus NRs as previously annotated^33^.

**Extended Data Figure 11**. (A) UMAP embeddings of immune cells as described in Fig. 3b visualized by responder status, IL6, IL8 or CLEC4D expression levels. (B) Correlation of IL8, VEGFA and CLEC4D normalized plasma protein expression with absolute monocyte count (AMC) or absolute neutrophil count (ANC) for each sample.

**Extended Data Figure 12**. Scatter plot illustrating the correlation of the difference between Rs and NRs of plasma protein expression compared to tumor immune cell gene expression from scRNAseq data.

**Extended Data Figure 13**. (A) Heatmap showing gene expression of non-response module proteins in single immune cells as described in Fig. 3b. (B) Dot plot showing average expression and percentage of cells expressing in non-response module protein in each cluster of immune cells as described in Fig. 3b.

**Extended Data Figure 14**. (A) UMAP embedding of 7 tumor epithelial cell samples from patients in this plasma proteomic cohort. Cells are labeled by sample ID. (B) Heatmap showing gene expression of non-response module proteins in single tumor epithelial cells as described in (A). (C) Non-response module enrichment score for each cell in (A) visualized on a UMAP embedding.

**Extended Data Figure 15**. (A) Heatmap of marker genes for monocyte and macrophage subclusters from data in Fig. 3b. (B) Heatmap showing gene expression of non-response module proteins in single monocytes or macrophages as described in (A). (C) Histograms of non-response module enrichment scores in cells within each of the monocyte and macrophage clusters described in (A). (D) Correlation between gene expression signatures of macrophage subsets in this study, and two other studies by Mulder et al.^38^ and Casanova-Acebes et al.^36^

**Extended Data Figure 16**. (A) Heatmap showing gene expression of non-response module proteins in single immune cells from an independent dataset^34,35^. (B)-(D) UMAP embedding of immune cell scRNAseq data from an independent dataset with annotation of (B) major cell types and subsets, (C) pre- or post-treatment designation and (D) non-response module enrichment score. (E) Heatmap showing gene expression of non-response module proteins in single tumor epithelial cells from an independent dataset. (F)-(G) UMAP embedding of tumor epithelial cell scRNAseq data from an independent dataset with annotation of (F) pre- or post-treatment designation and (G) non-response module enrichment score.

**Extended Data Figure 17**. Scatter plot showing gene expression counts as log(TPM+1), for all response-effect proteins discovered in plasma, in tumor epithelial cells versus immune within the TME. Data represented is obtained by scRNAseq analysis of biopsies obtained from patients in this plasma proteomic cohort either from previously generated immune cell subsets^33^ or newly generated tumor epithelial cells.

**Extended Data Figure 18**. Scatter plot showing gene expression counts as log(TPM+1) from an independent dataset^34,35^, for all response-effect proteins discovered in plasma, in tumor epithelial cells versus (A) tumor immune cells and (B) tumor macrophages within the TME.

**Extended Data Figure 19**. Analysis of bulk RNA-sequencing data from an independent cohort for prognostic implications of the non-response module^8^. (A) Correlation between non-response module scores and myeloid scores (patient samples n = 103, Spearman’s rank correlation coefficient, R= 0.65, p<0.001). (B) Distribution of myeloid scores, separated by Rs (patient samples n = 47) and progressors (n= 56). Median of myeloid scores of both Rs and progressors (n = 103) is shown. (C) Overall survival (OS) and (D) progression-free survival (PFS) stratified by high versus low non-response module score (split by median) in a subset of patients from Schadendorf cohort with relatively high myeloid proportions (n= 60). Tumors with high non-response module trended towards worse OS and PFS (two-sided KM log-rank test,n.s.). (E) Overall survival (OS) stratified by quartiles of high versus low non-response module score in a subset of patients from Schadendorf cohort with relatively high myeloid proportions (n= 60). Tumors with high non-response module had worse OS (top quartile, two-sided KM log-rank test, P = 0.023), with consistent trend in PFS (two-sided KM log-rank test, n.s.).

## Supplemental Table legends

**Table S1**. Comprehensive list of Olink panels and proteins used in this study.

**Table S2**. Summary table of characteristics of patients in the discovery cohort of this study.

**Table S3**. Results from linear-mixed effect models and post-hoc testing for plasma proteomic data to identify differential proteins for time, response and interaction effect.

**Table S4**. Proteins in the plasma non-response module.

**Table S5**. Differential marker genes for myeloid clusters from single-cell RNA-sequencing data.

