## Supplemental Figures for "Plasma proteomic biomarkers identify non-responders and reveal biological insights about the tumor microenvironment in melanoma patients after PD1 blockade"

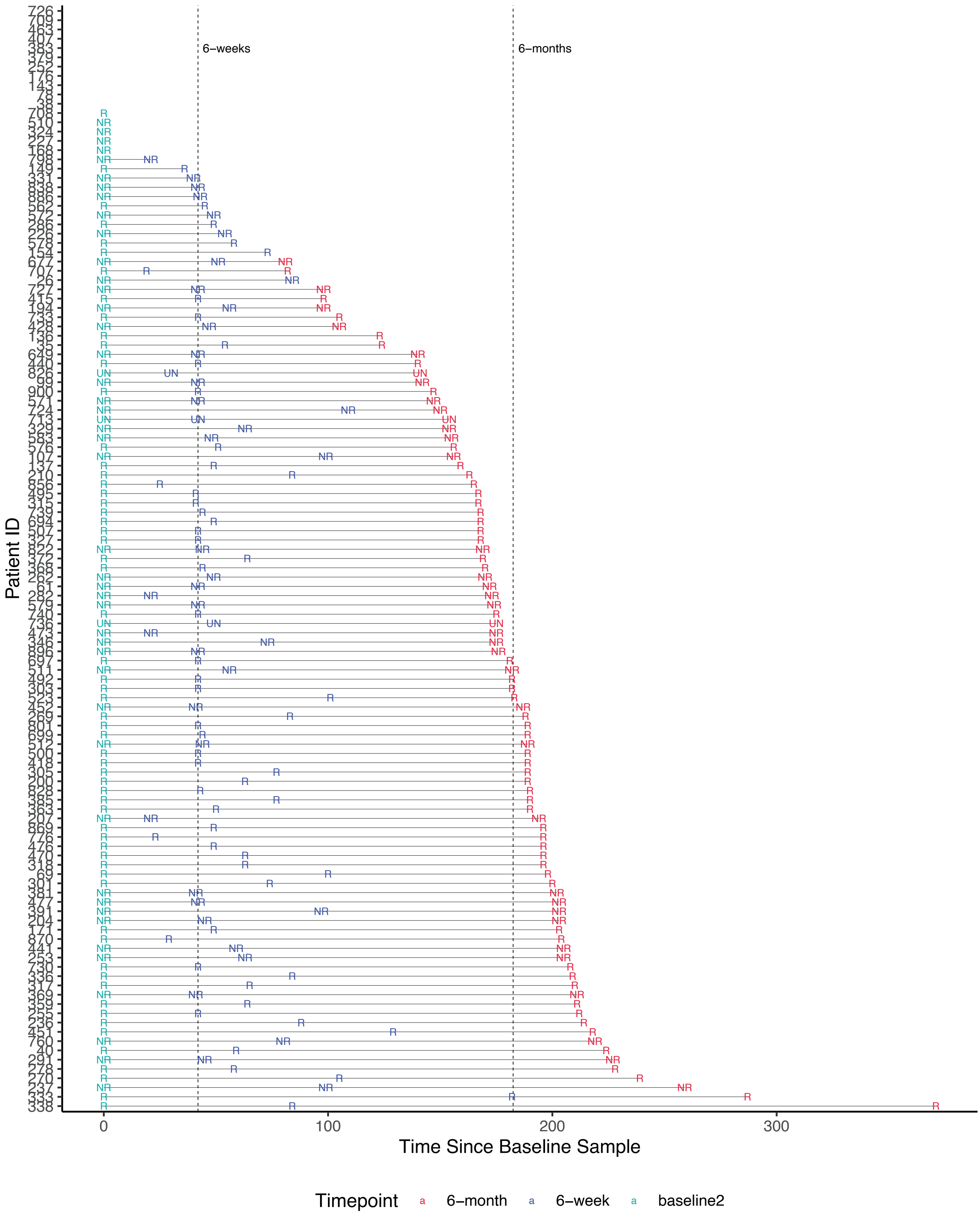


### Extended Data Figure 1

A

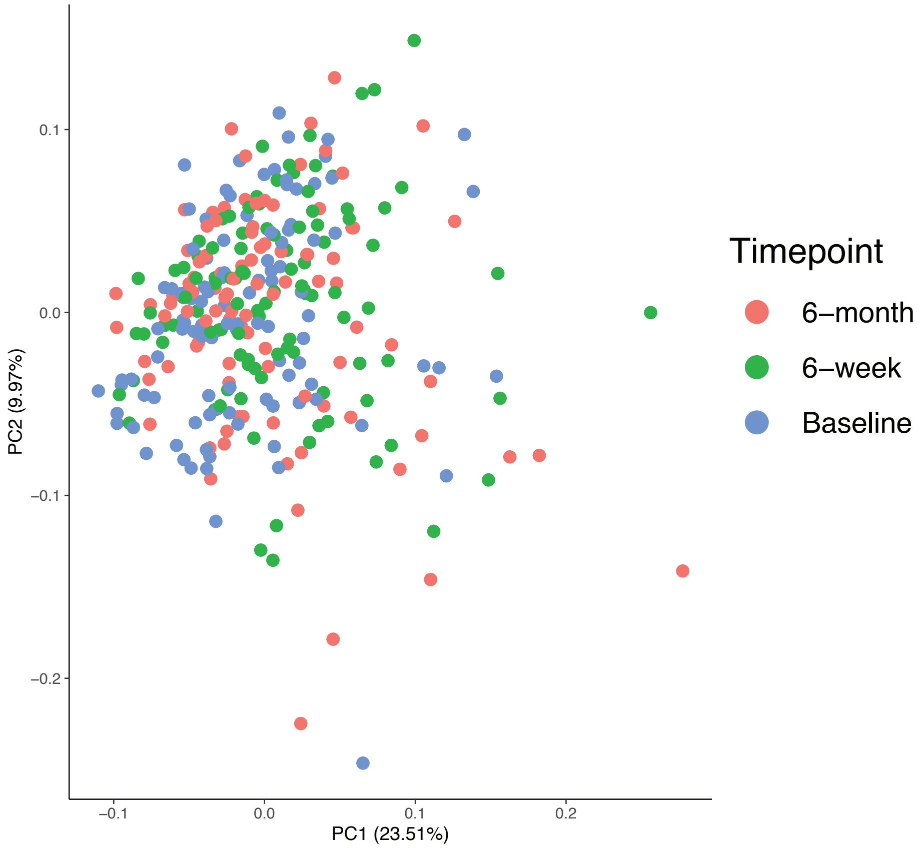


B

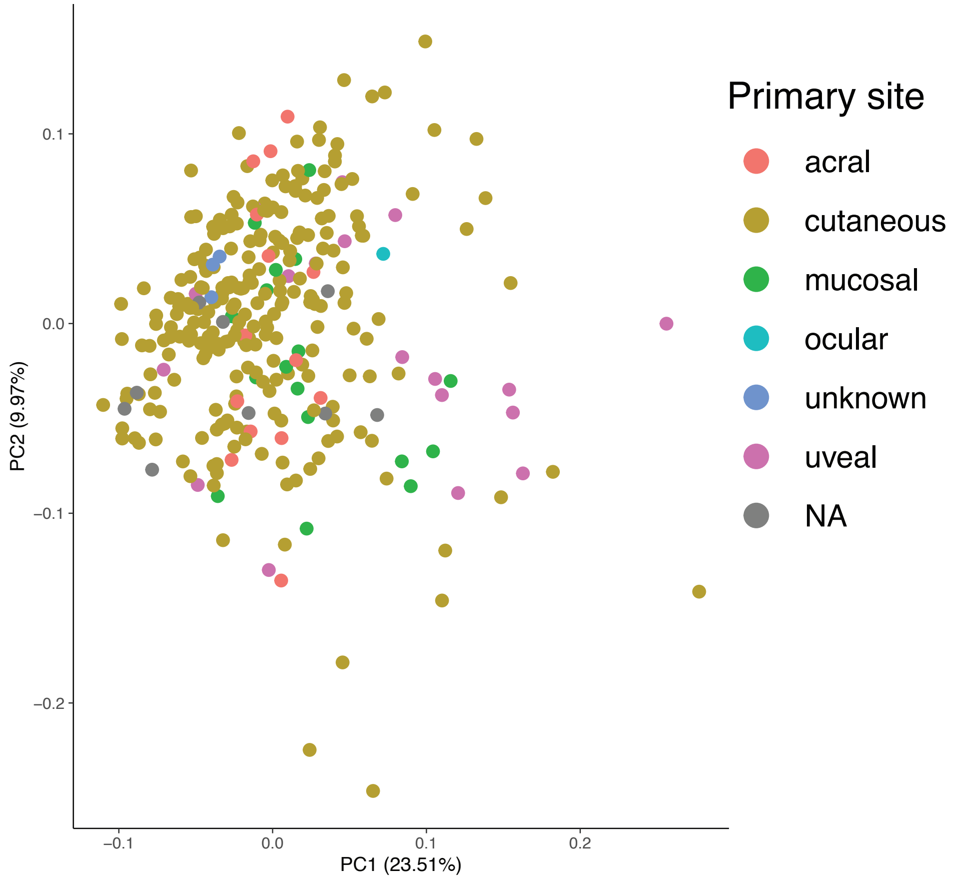


**Extended Data Figure 2**

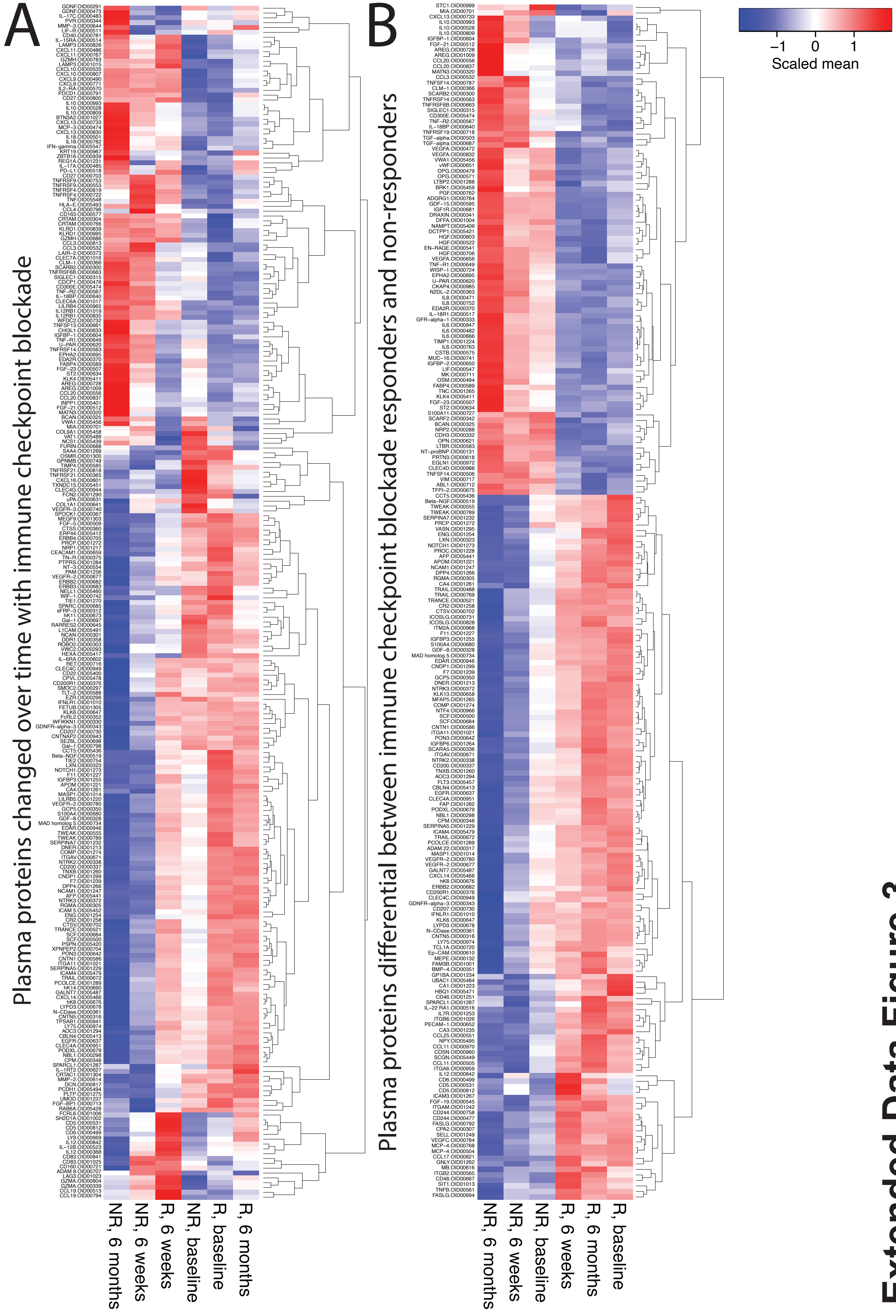


Extended Data Figure 3

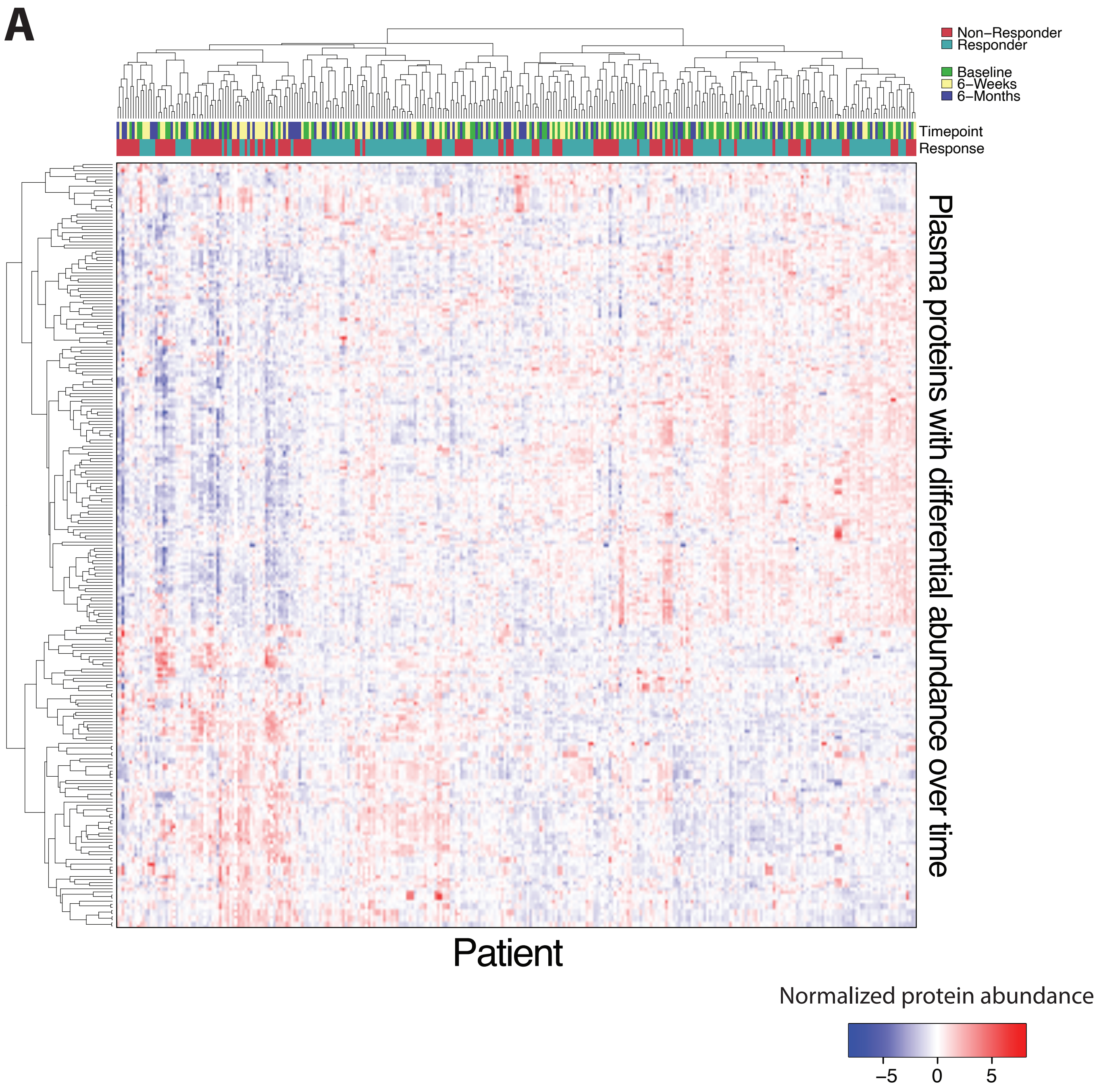


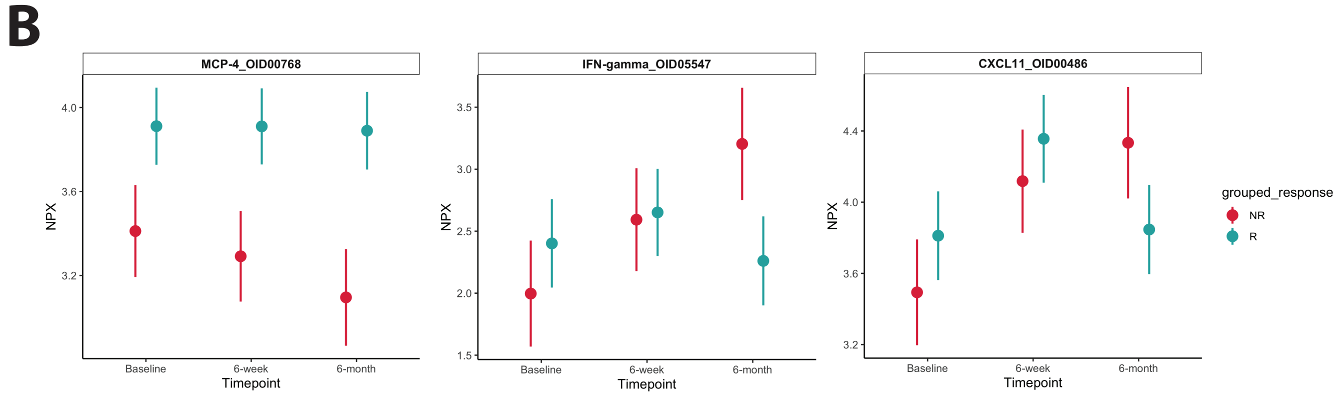


### Extended Data Figure 4

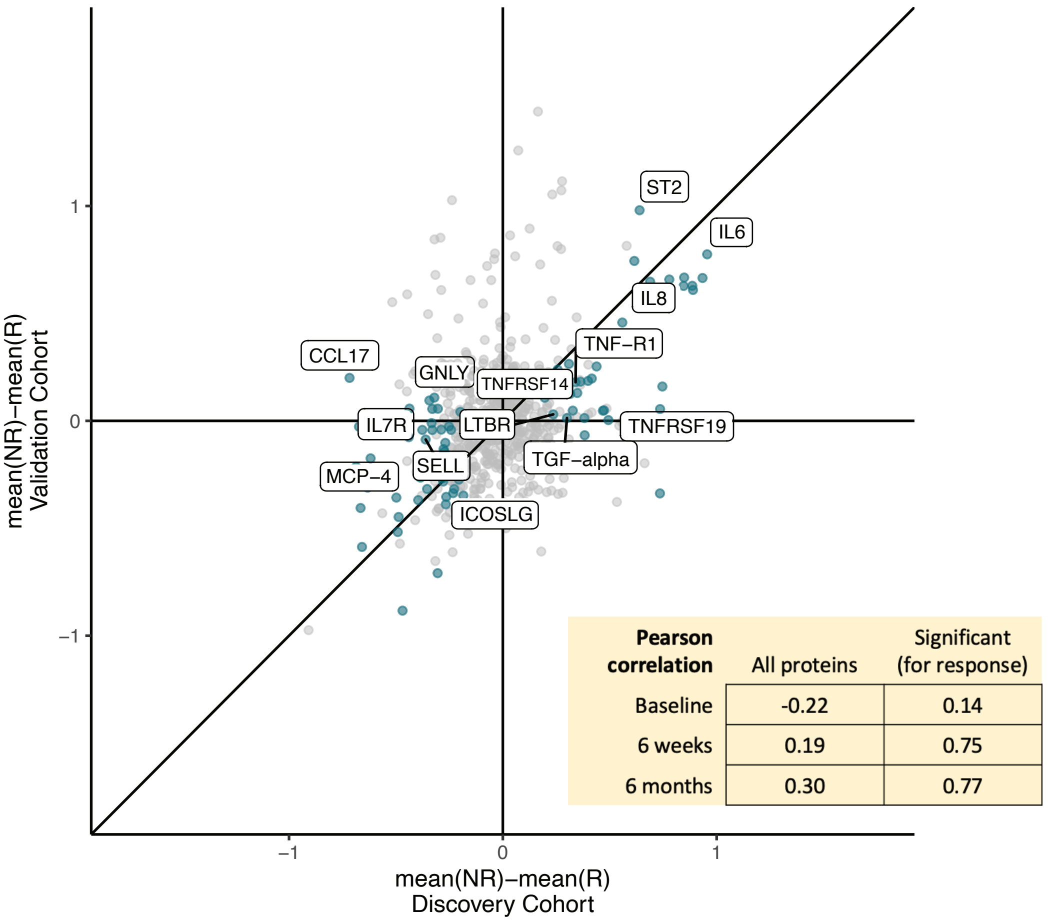


| Pearson correlation | All proteins | Significant (for response) |
|---|---|---|
| Baseline | -0.22 | 0.14 |
| 6 weeks | 0.19 | 0.75 |
| 6 months | 0.30 | 0.77 |

### Extended Data Figure 5

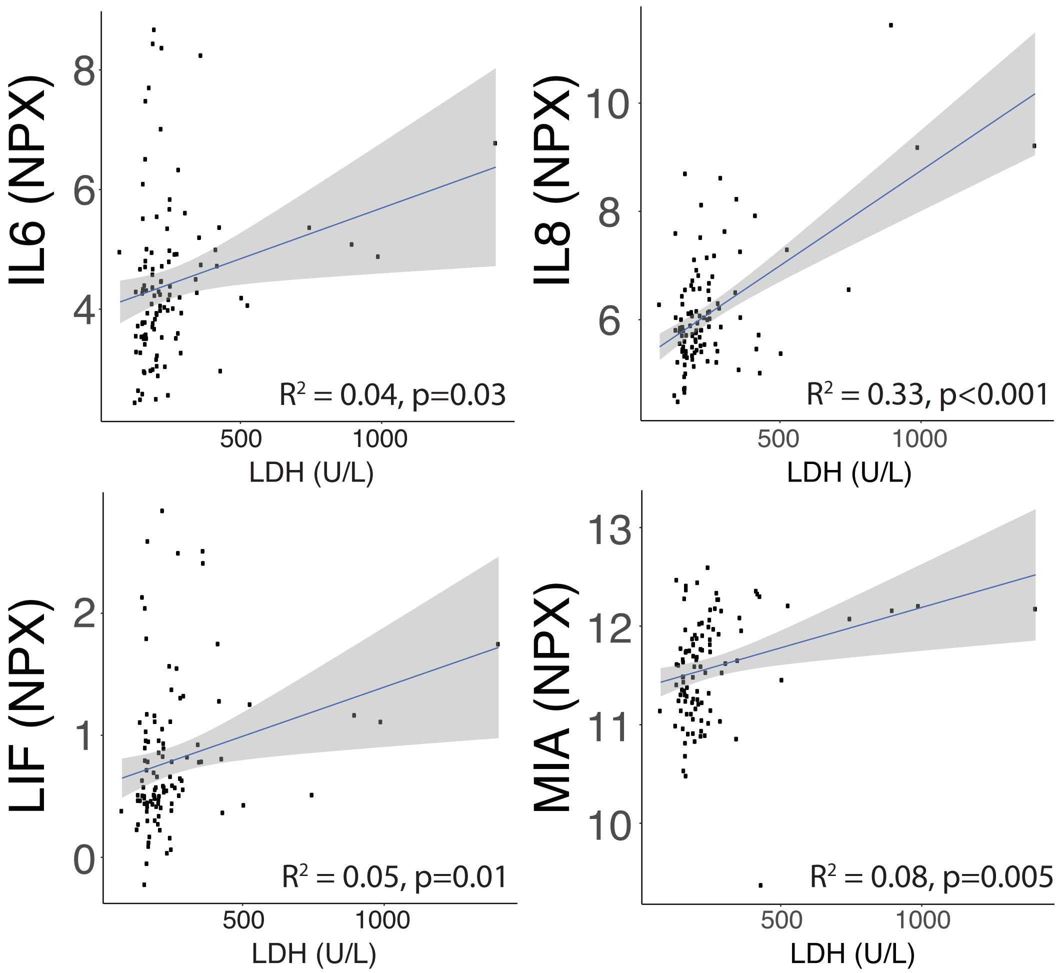


### Extended Data Figure 6

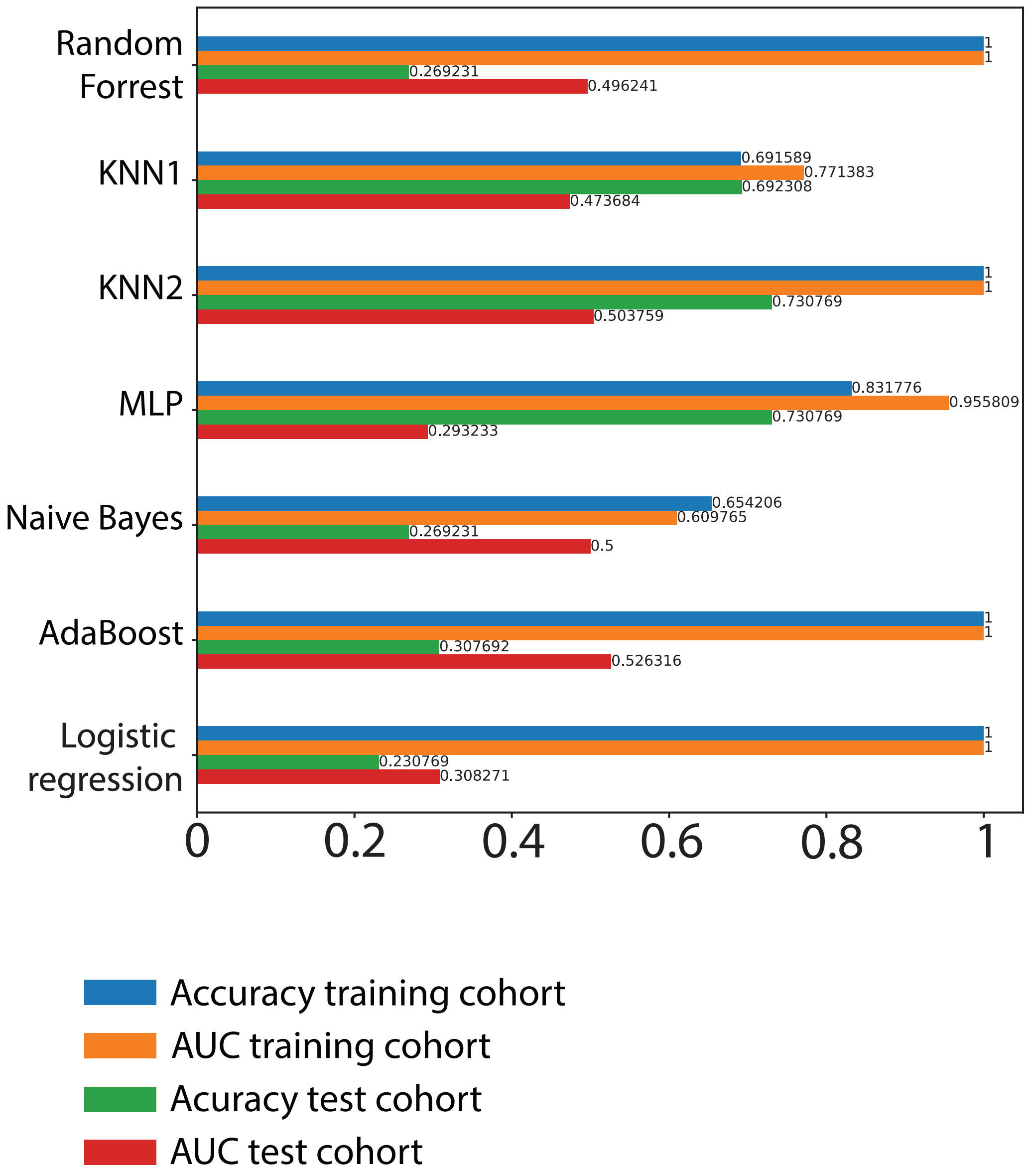


**Extended Data Figure 7**

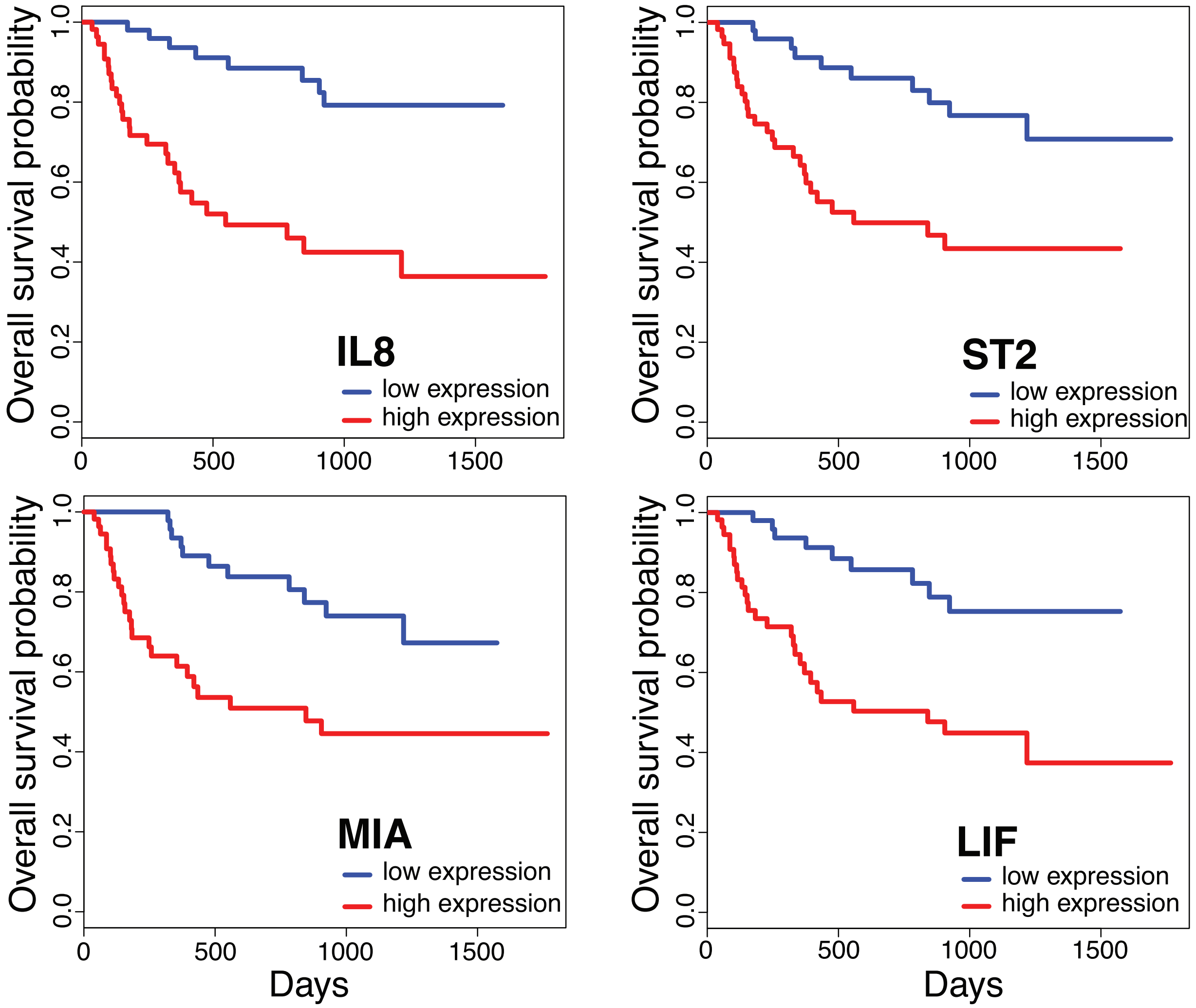


**Extended Data Figure 8**

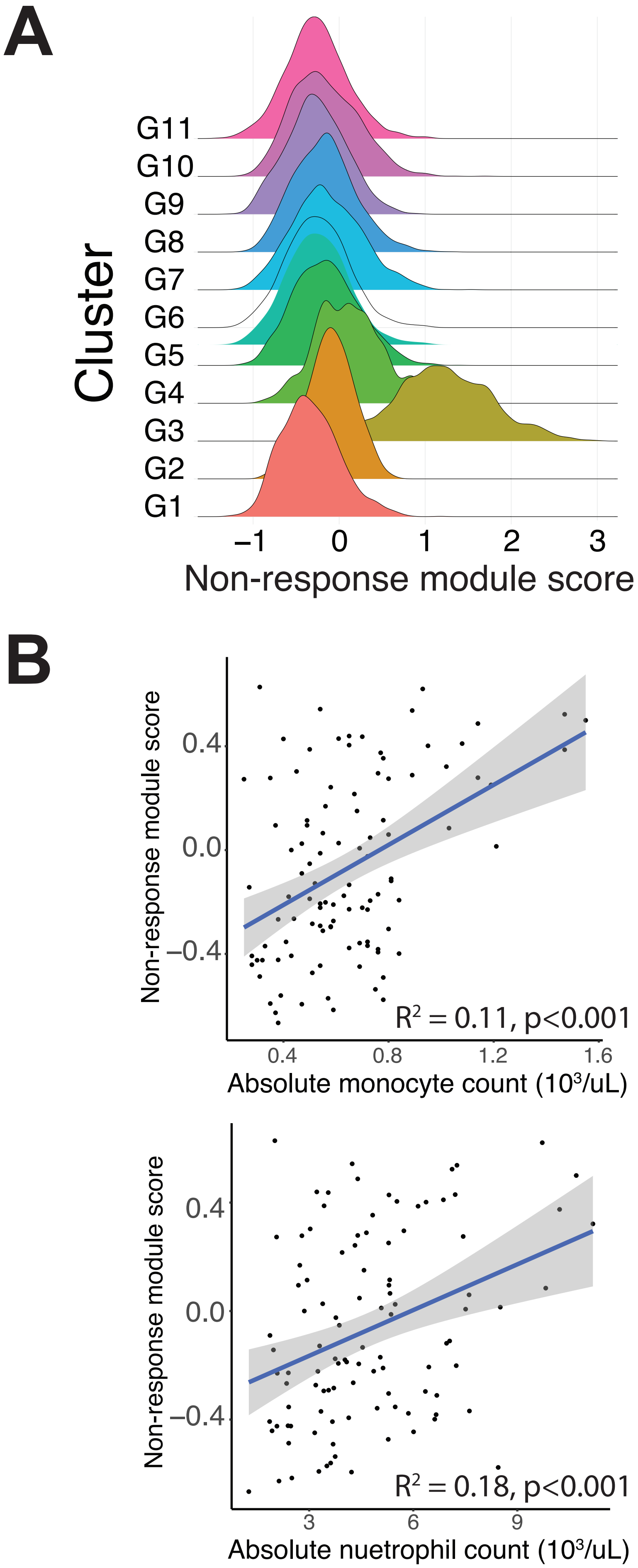


**Extended Data Figure 9**

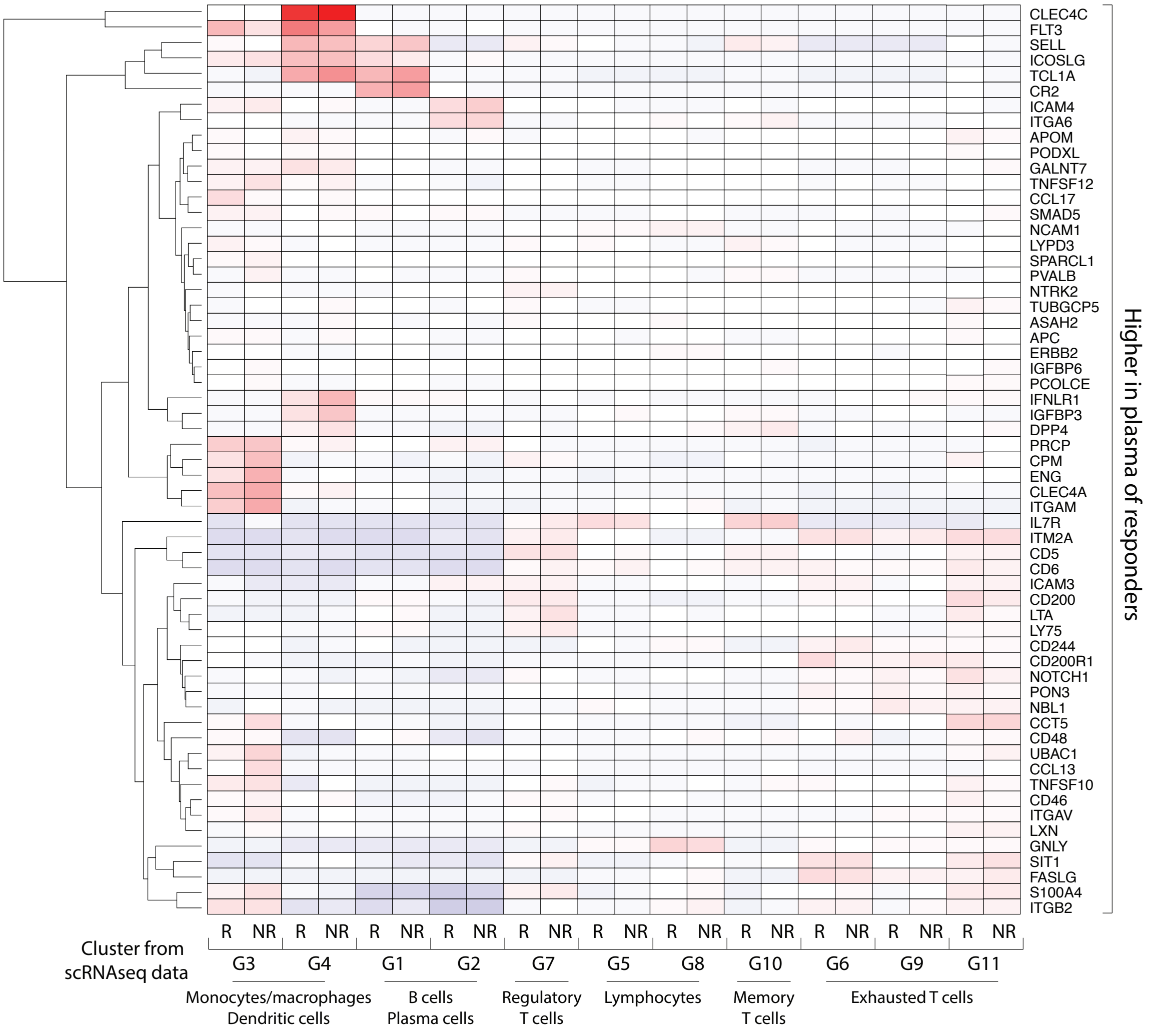


### Extended Data Figure 10

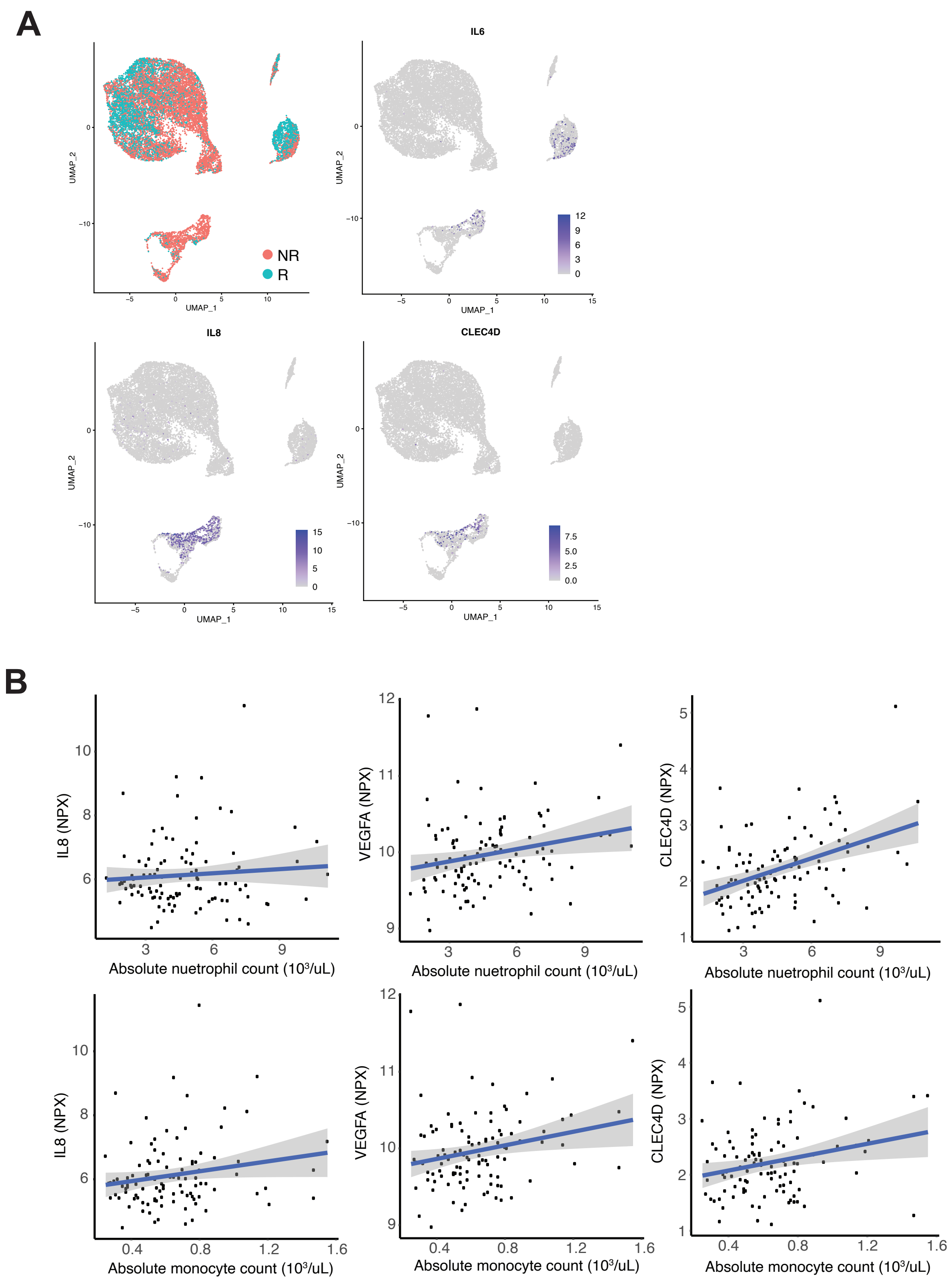


### Extended Data Figure 11

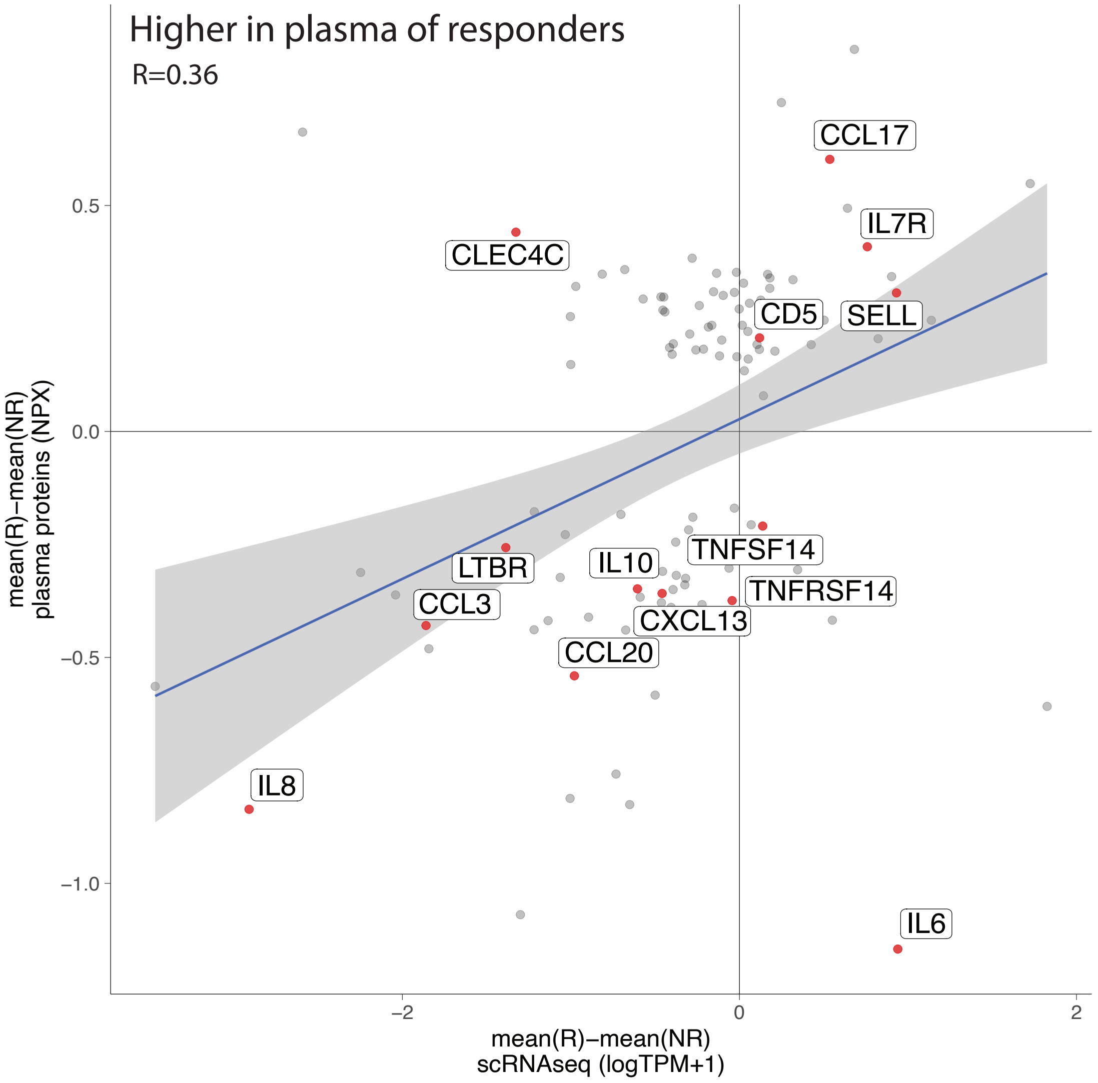


**Extended Data Figure 12**

A

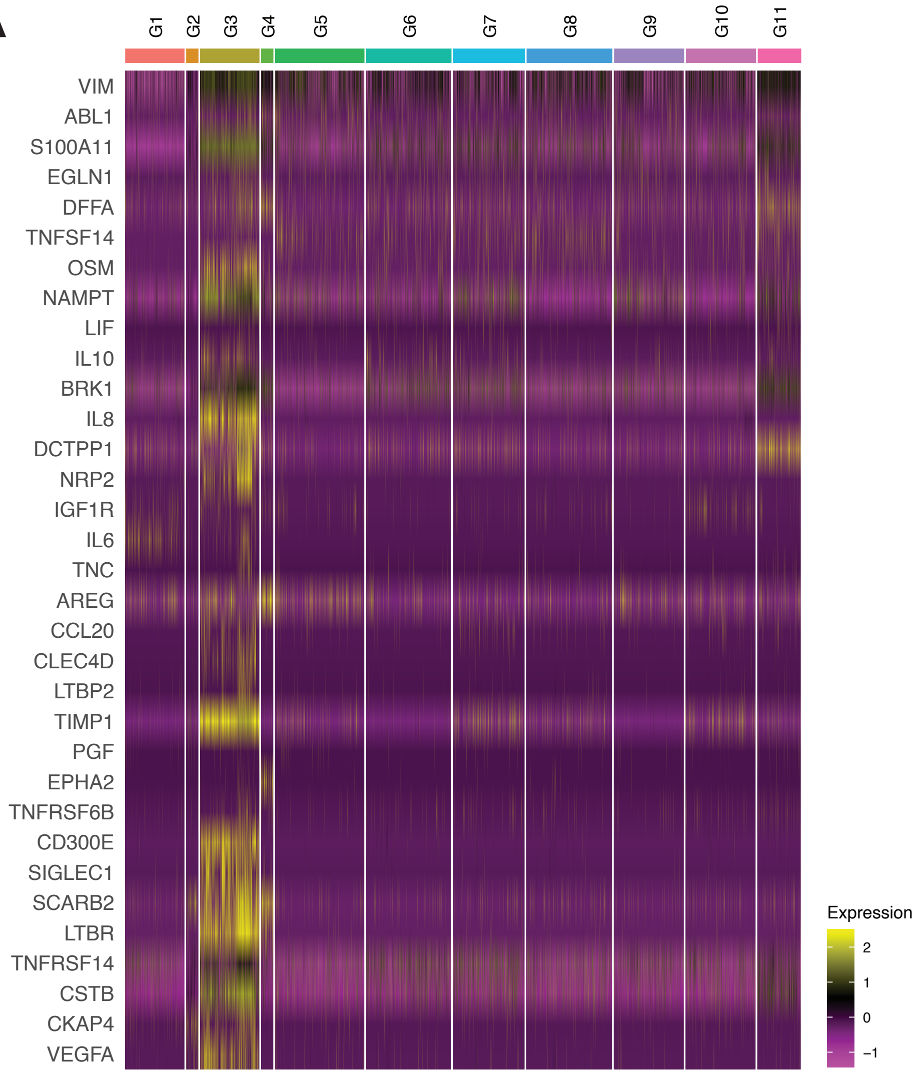


B

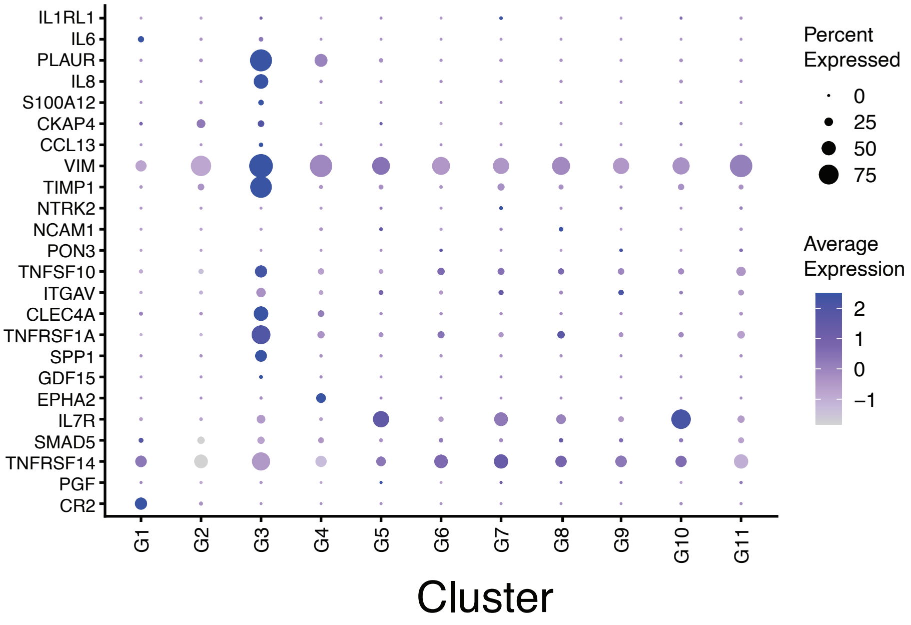


### Extended Data Figure 13

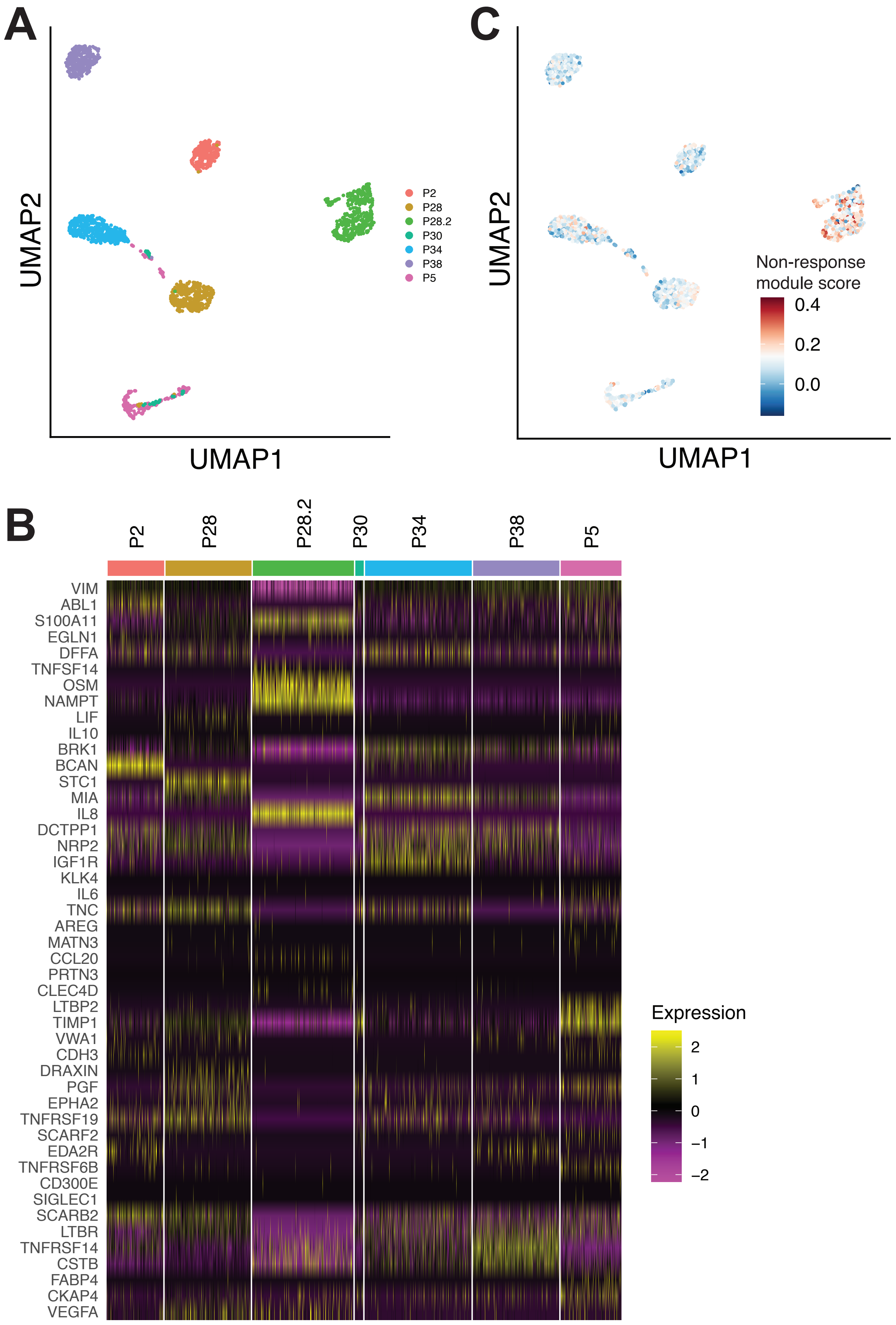


**Extended Data Figure 14**

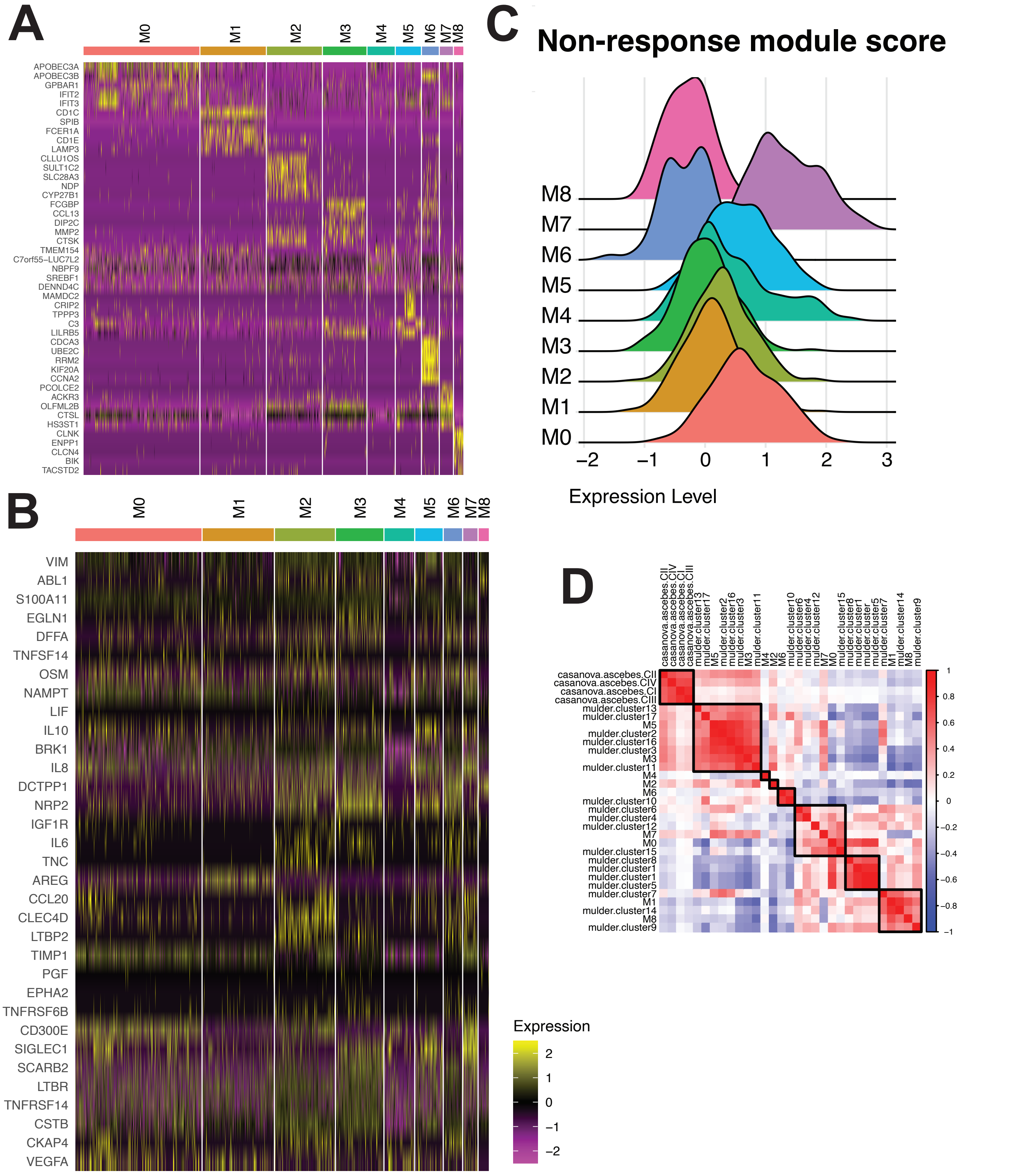


### Extended Data Figure 15

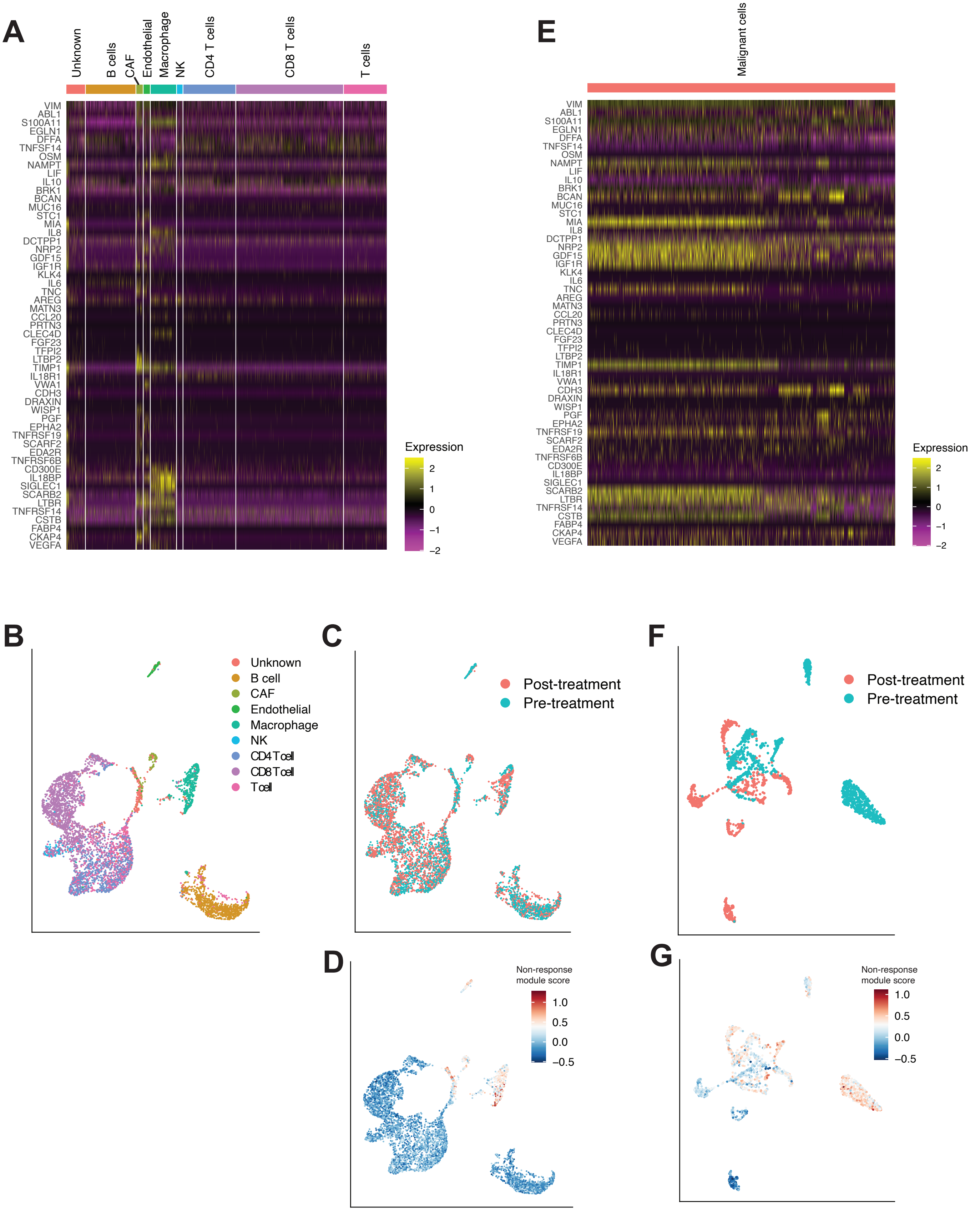


### Extended Data Figure 16

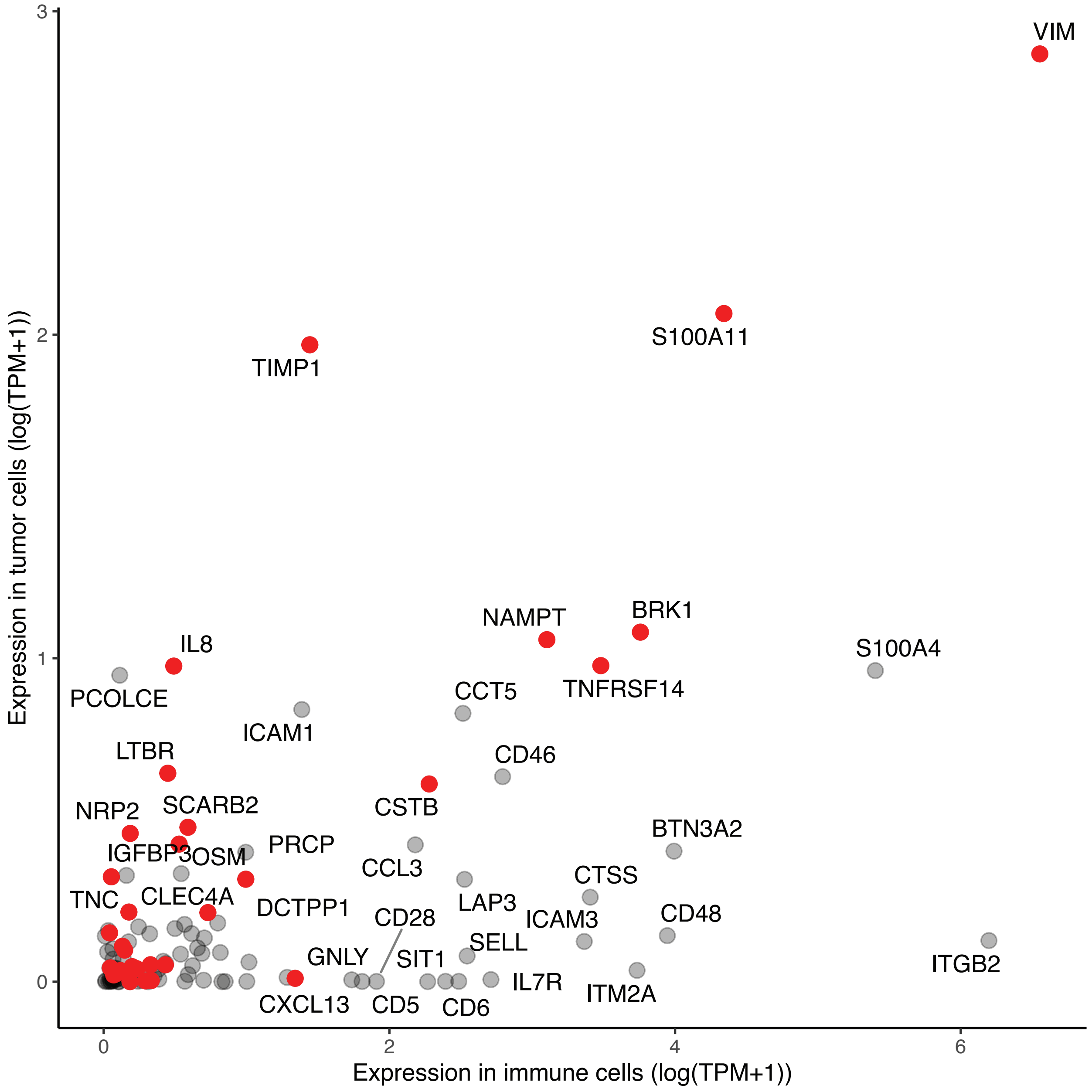


### Extended Data Figure 17

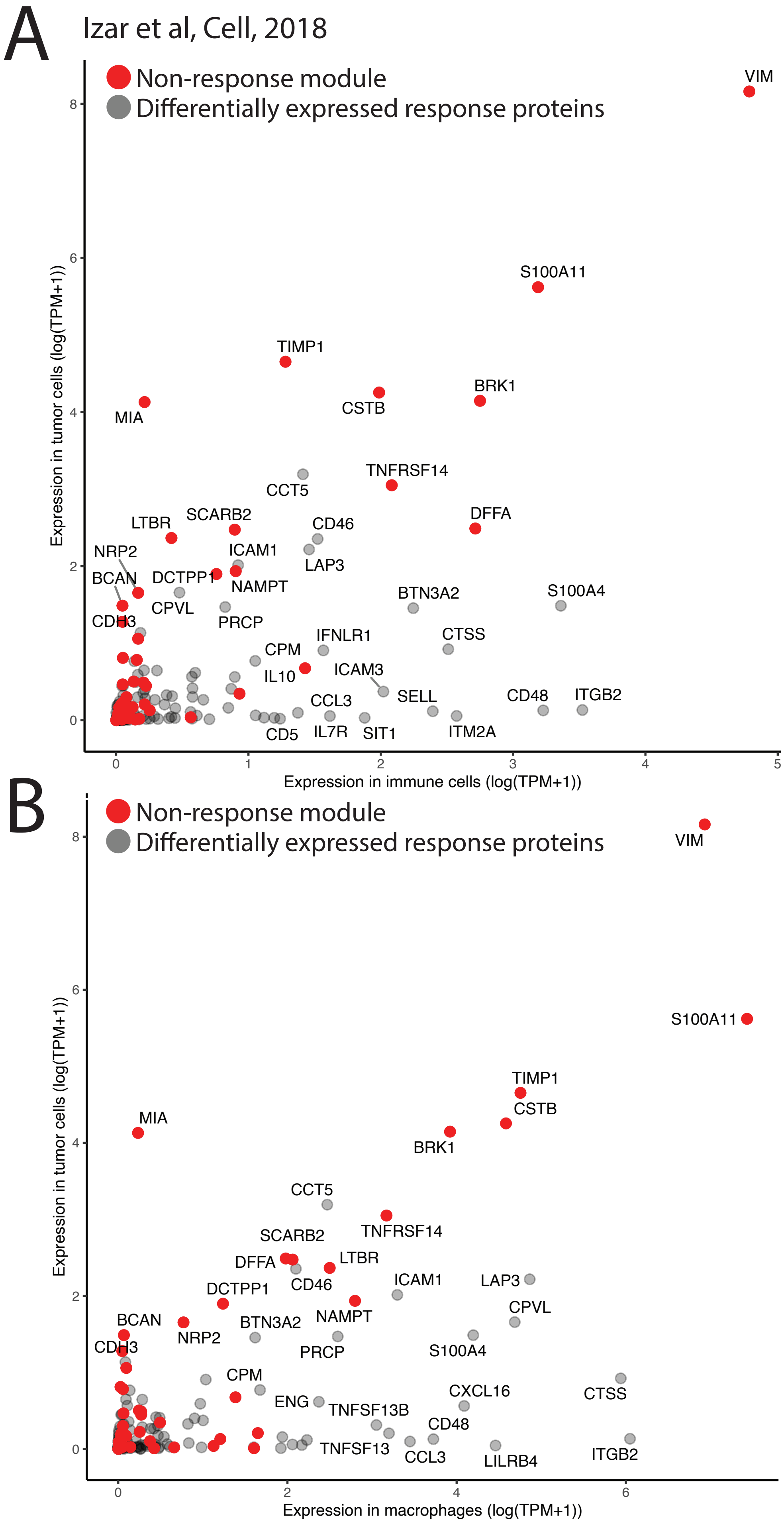


**Extended Data Figure 18**

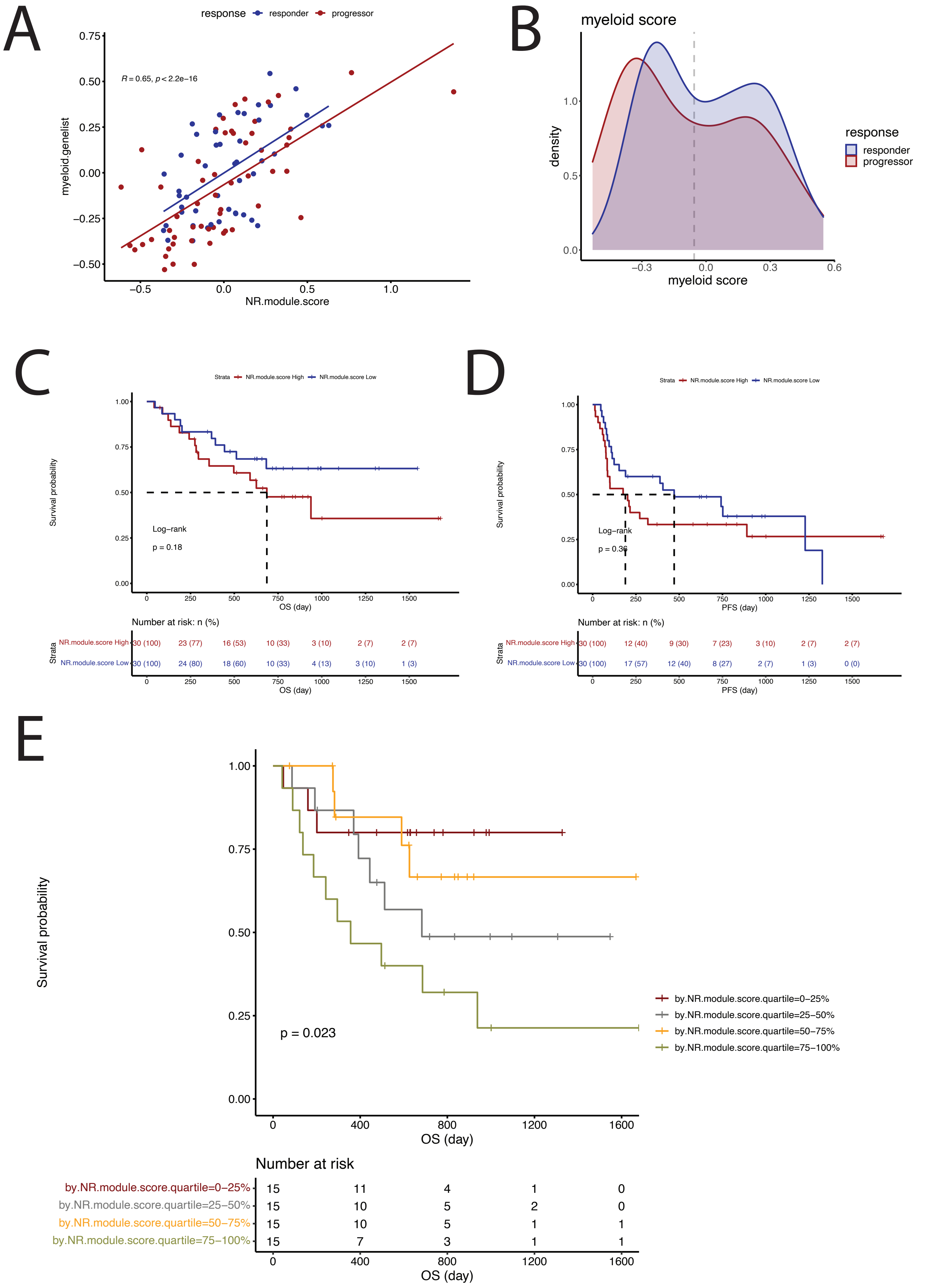


**Extended Data Figure 19**
