## Supplemental Table 2 for "Plasma proteomic biomarkers identify non-responders and reveal biological insights about the tumor microenvironment in melanoma patients after PD1 blockade"

**Table S2**: Summary table of characteristics of patients in the discovery cohort of this study.

| | All patients | Responders | Non-responders |
|---|---|---|---|
| **Total patients** | 116 | 66 | 50 |
| Age, mean (range) | 62 (25-97) | 61 (25-90) | 63 (30-97) |
| Sex, # females (%) | 37 (32%) | 17 (26%) | 19 (38%) |
| **Primary site, N (%)** | | | |
| Cutaneous | 93 (80%) | 60 (90%) | 33 (66%) |
| Acral | 5 (4%) | 1 (2%) | 4 (8%) |
| Ocular | 8 (7%) | 2 (3%) | 6 (12%) |
| Mucosal | 8 (7%) | 2 (3%) | 6 (12%) |
| Unknown | 2 (2%) | 1 (2%) | 1 (2%) |
| **Stage, N (%)** | | | |
| M0 | 8 (7%) | 5 (8%) | 3 (6%) |
| M1a | 11 (9%) | 6 (9%) | 5 (10%) |
| M1b | 21 (18%) | 14 (21%) | 7 (14%) |
| M1c | 46 (40%) | 25 (38%) | 21 (42%) |
| M1d | 29 (25%) | 16 (24%) | 13 (26%) |
| Unknown | 1 (1%) | 0 (0%) | 1 (2%) |
| **Brain metastases, N (%)** | 48 (41%) | 23 (34%) | 25 (50%) |
| **ECOG PS, N (%)** | | | |
| 0 | 62 (53%) | 37 (56%) | 25 (50%) |
| 1 | 36 (31%) | 21 (32%) | 15 (30%) |
| 2/3 | 8 (7%) | 2 (3%) | 6 (12%) |
| Unknown | 10 (9%) | 6 (9%) | 4 (8%) |
| **Treatment, N (%)** | | | |
| aPD1 / aPDL1 | 81 (70%) | 43 (65%) | 38 (76%) |
| aCTLA4 | 10 (9%) | 4 (6%) | 6 (12%) |
| aPD1 + aCTLA4 | 25 (22%) | 19 (29%) | 6 (12%) |
| **Labs, mean (range)** | | | |
| LDH (U/L) | 249 (46-1404) | 212 (46-527) | 298 (103-1404) |
| White blood count ($10^3$/uL) | 7.3 (3.0-21.6) | 6.9 (3.0-16.8) | 7.9 (3.3-21.6) |
| Hemoglobin | 13.2 (9.2-16.4) | 13.5 (9.3-16.4) | 12.9 (9.2-16.2) |
| Absolute neutrophil count ($10^3$/uL) | 4.8 (1.3-18.6) | 4.3 (1.3-8.5) | 5.4 (1.9-18.6) |
| Absolute monocyte count ($10^3$/uL) | 0.6 (0.3-1.6) | 0.6 (0.3-1.5) | 0.7 (0.3-1.6) |
| Absolute lymphocyte count ($10^3$/uL) | 1.6 (0.4-8.5) | 1.6 (0.5-8.5) | 1.5 (0.4-3.6) |
